# *In vitro* evolution of colistin resistance in the *Klebsiella pneumoniae* complex follows multiple evolutionary trajectories with variable effects on fitness and virulence characteristics

**DOI:** 10.1101/2020.05.24.112334

**Authors:** Axel B. Janssen, Dennis J. Doorduijn, Grant Mills, Malbert R.C. Rogers, Marc J.M. Bonten, Suzan H.M. Rooijakkers, Rob J.L. Willems, Jose A. Bengoechea, Willem van Schaik

**Author notes:** Address correspondence to Willem van Schaik,.

## Abstract

The increasing prevalence of multidrug-resistant Gram-negative opportunistic pathogens, including *Klebsiella pneumoniae*, has led to a resurgence in the use of colistin as a last-resort drug. Colistin is a cationic lipopeptide antibiotic that selectively acts on Gram-negative bacteria through electrostatic interactions with anionic phosphate groups of the lipid A moiety of lipopolysaccharides (LPS). Colistin resistance in *K. pneumoniae* is mediated through loss of these phosphate groups, or modification with cationic groups (e.g. 4-amino-4-deoxy-L-arabinose (L-Ara4N), or phosphoethanolamine), but also hydroxylation of acyl-groups of lipid A. Here, we study the *in vitro* evolutionary trajectories towards colistin resistance in clinical *K. pneumoniae* complex strains (three *K. pneumoniae sensu stricto* strains and one *K. variicola* subsp. *variicola* strain) and their impact on fitness and virulence characteristics.

Through population sequencing during the *in vitro* evolution experiment, we found that resistance develops through a combination of single nucleotide polymorphisms (SNPs), insertion and deletions (indels), and the integration of insertion sequence (IS) elements, affecting genes associated with LPS biosynthesis and modification, and capsule structures. The development of colistin resistance decreased the maximum growth rate of one *K. pneumoniae sensu stricto* strain, but not in the other three *K. pneumoniae sensu lato* strains. Colistin-resistant strains had lipid A modified through hydroxylation, palmitoylation, and L-Ara4N addition. Colistin-resistant *K. pneumoniae sensu stricto* strains exhibited cross-resistance to LL-37, in contrast to the *K. variicola* subsp. *variicola* strain that did not change in susceptibility to LL-37. Virulence, as determined in a *Caenorhabditis elegans* survival assay, was higher in two colistin-resistant strains.

Our study suggests that nosocomial *K. pneumoniae* complex strains can rapidly develop colistin resistance *de novo* through diverse evolutionary trajectories upon exposure to colistin. This effectively shortens the lifespan of this last-resort antibiotic for the treatment of infections with multidrug-resistant *Klebsiella*.

**Author summary:** Bacteria that frequently cause infections in hospitalised patients are becoming increasingly resistant to antibiotics. Colistin is a positively charged antibiotic that is used for the treatment of infections with multidrug-resistant Gram-negative bacteria. Colistin acts by specifically interacting with the negatively charged LPS molecule in the outer membrane of Gram-negative bacteria. Colistin resistance is mostly mediated through modification of LPS to reduce its negative charge. Here, we use a laboratory evolution experiment to show that strains belonging to the *Klebsiella pneumoniae* complex, a common cause of multidrug-resistant hospital-acquired infections, can rapidly accumulate mutations that reduce the negative charge of LPS without an appreciable loss of fitness. Colistin resistance can lead to cross-resistance to an antimicrobial peptide of the human innate immune system, but can increase susceptibility to serum, and virulence in a nematode model. These findings show that extensively resistant *K. pneumoniae* complex strains may rapidly develop resistance to the last-resort antibiotic colistin via different evolutionary trajectories, while retaining their ability to cause infections.

## Introduction

*Klebsiella pneumoniae* is a Gram-negative opportunistic pathogen and a leading cause of hospital-associated infections such as pneumonia, soft tissue infections, and urinary tract infections. *K. pneumoniae* may also asymptomatically colonize the skin, upper respiratory tract, and digestive tract of healthy individuals [1,2]. The *K. pneumoniae* complex is genetically diverse, with different phylogroups within the complex corresponding to different species and sub-species, each occupying specific niches [1,2]. The *K. pneumoniae sensu strico* and *K. quasipneumoniae* phylogroups are associated with human intestinal carriage, whilst the *K. variicola* phylogroup is associated with plants and bovine carriage [1,3]. Of all strains isolated from human infections and typed as *K. pneumoniae*, the majority is *K. pneumoniae sensu stricto*, but *K. variicola* and *K. quasipneumoniae* have also been found to cause infections in patients and are frequently misidentified as *K. pneumoniae* [4,5]. Although infections with strains from the *K. variicola* phylogroup are relatively rare, they have been associated with the highest mortality rate within the *K. pneumoniae* complex [3].

In recent years, *K. pneumoniae* complex strains have rapidly emerged as multidrug-resistant pathogens through acquisition of resistance to third-generation cephalosporins, fluoroquinolones, and aminoglycosides, and have increasingly become resistant to carbapenems through the acquisition of carbapenemases [6–9]. The increasing prevalence of multidrug resistance within the *K. pneumoniae* complex, and the lack of development of novel antibiotic classes effective against Gram-negative bacteria, have limited the available therapeutic options against multidrug-resistant *K. pneumoniae* complex strains. These limitations have prompted the resurgence in the use of the antibiotic colistin in treatment of infections by *K. pneumoniae* complex strains [10–13]. After its introduction into clinical practice in the 1950s, colistin fell into disuse in human medicine in the 1970s because of the neuro- and nephrotoxic side effects associated with its use, and the development of safer classes of antibiotics. Due to the emergence multidrug-resistant Gram-negative opportunistic pathogens, like *K. pneumoniae*, it has recently regained clinical relevance as a last-line antibiotic [13–16].

Colistin (polymyxin E) is a cationic, amphipathic molecule composed of a fatty acid chain linked to a non-ribosomally synthesized decapeptide [17,18]. The mechanism of action of colistin relies on the selective presence of the negatively charged lipopolysaccharides (LPS) in the membranes of Gram-negative bacteria. The negative charges of LPS are carried by the anionic phosphate groups of the lipid A moiety of LPS, which enable colistin to bind through electrostatic interactions [17–20]. Insertion of colistin into the outer membrane leads to membrane permeabilization. The subsequent destabilization of the cytoplasmic membrane, where LPS is present after synthesis in the cytoplasm while awaiting transport to the outer membrane, ultimately leads to cell death [17,19,21,22].

The increased use of colistin to treat infections with multidrug-resistant Gram-negative bacteria, especially in low- and middle-income countries [13], and the use of colistin in livestock farming, either therapeutically to treat enteric infections or as a growth promoter [23,24], has led to a rise in colistin resistance in *K. pneumoniae* from clinical, veterinary, and environmental sources [9,25–29]. Colistin resistance in *K. pneumoniae* complex strains is mostly mediated through decoration of lipid A with cationic groups, to counteract the electrostatic interactions between colistin and lipid A [17]. These modifications can be the result of point mutations and indels in chromosomally located genes (including *phoPQ*, *pmrAB*, and *crrAB*) resulting in amino acid substitutions and frameshift mutations, respectively. In addition, the acquisition of mobile genetic elements carrying a member of the *mcr*-gene family may also lead to lipid A modification [30–35]. In *K. pneumoniae*, the inactivation of *mgrB* encoding an negative regulator of PhoPQ, through the insertion of an insertion sequence (IS) element, or a mutation leading to the formation of a premature stop codon, is a particularly frequently observed colistin resistance mechanism [25,36–41]. Other mechanisms of colistin resistance in *K. pneumoniae* include the upregulated expression of efflux pumps [42,43], changes in LPS production [25,44], and the overproduction of capsular polysaccharides [45,46].

Upon infection the innate immune system will attempt to neutralize invading bacteria. The cellular components of the innate immune system can detect Gram-negative bacteria through the presence of LPS [47]. Activated immune cells can kill bacteria and will attempt to kill them by unleashing bactericidal components including the antimicrobial peptide LL-37. Similar to colistin, LL-37 relies on electrostatic interactions with LPS for its mechanism of action [48]. Modifications to LPS may influence the efficacy of bactericidal components, and may thus result in altered virulence by reducing the effectiveness of these components [47–51]. Modifications capable of affecting the efficiency of the immune system include neutralization of the anionic charges carried by lipid A, and changes in acylation of lipid A [47,49,50,52]. These changes are mediated through the PhoPQ and PmrAB two-component regulatory systems. Notably, colistin resistance is mediated through the same modifications and two-component regulatory systems. The development of colistin resistance may thus also affect virulence characteristics.

To better understand the mechanisms and consequences of colistin resistance in *K. pneumoniae* complex strains, we determined the evolutionary trajectories of three *K. pneumoniae sensu stricto* strains and one *K. variicola* subsp. *variicola* strain towards colistin resistance in an *in vitro* evolution experiment, and determined how colistin resistance impacted fitness, LPS modifications, and virulence characteristics.

## Materials and Methods

### Ethical statement

The colistin-susceptible *K. pneumoniae* complex strains used in this study were isolated as part of routine diagnostic procedures, which did not require consent or ethical approval by an institutional review board.

### Bacterial strains, growth conditions, and chemicals

The colistin-susceptible KP209, KP040, KP257, and KV402 strains were retrospectively, obtained from the diagnostic laboratory of the University Medical Center Utrecht in Utrecht, the Netherlands. In initial routine diagnostic procedures, they were identified as *K. pneumoniae sensu stricto* by matrix-assisted laser desorption–ionisation time-of-flight (MALDI-TOF) on a Bruker microflex system (Leiderdorp, The Netherlands). Colistin susceptibility testing of the clinical isolates was initially performed on a BD Phoenix automated identification and susceptibility testing system (Becton Dickinson, Vianen, The Netherlands). All strains were grown either in lysogeny broth (LB; Oxoid, Landsmeer, The Netherlands) with agitation at 300 rpm, or on LB agar, at 37°C, unless otherwise specified. Colistin sulphate was obtained from Duchefa Biochemie (Haarlem, The Netherlands).

### Determination of minimal inhibitory concentration of colistin

Minimal inhibitory concentrations (MICs) to colistin were determined as described previously [53] in line with the recommendations from the joint Clinical & Laboratory Standards Institute and European Committee on Antimicrobial Susceptibility Testing (EUCAST) Polymyxin Breakpoints Working Group (http://www.eucast.org/fileadmin/src/media/PDFs/EUCAST_files/General_documents/Recommendations_for_MIC_determination_of_colistin_March_2016.pdf). In short, colistin susceptibility testing was performed using BBL™ Mueller Hinton II (cation-adjusted) broth (MHCAB; Becton Dickinson), untreated Nunc 96-wells round bottom polystyrene plates (Thermo Fisher Scientific, Landsmeer, The Netherlands), and Breathe-Easy sealing membranes (Sigma-Aldrich, Zwijndrecht, The Netherlands). The MIC was observed after stationary, overnight growth at 37°C, and was determined to be the lowest concentration where no visible growth was observed. The breakpoint value for colistin resistance of an MIC >2 µg/ml was obtained from EUCAST [54] (http://www.eucast.org/clinical_breakpoints/).

### *In vitro* evolution of colistin resistance

The nosocomial, colistin-susceptible *K. pneumoniae* strains were evolved towards colistin resistance by culturing in increasing colistin concentrations over a period of 5-7 days. Prior to the *in vitro* evolution experiments, MICs to colistin were determined in LB. Each strain was grown in 1 ml LB with initial colistin concentrations of 1 and 2 times the MIC. After overnight growth, 1 μl of the cultures with the highest concentration of colistin that had visible growth were used to propagate a fresh culture by inoculating 1 ml of fresh LB, supplemented with the same or twice the concentration of colistin in which growth was observed in the previous day’s culture (Supplemental Figure S1). This process was repeated for 5-7 days. Each overnight culture was stored at −80°C in 20% glycerol.

### Genomic DNA isolation and whole-genome sequencing

Genomic DNA was isolated using the Wizard Genomic DNA purification kit (Promega, Leiden, The Netherlands) according to the manufacturer’s instructions. DNA concentrations were measured with the Qubit 2.0 fluorometer and the Qubit dsDNA Broad Range Assay kit (Life Technologies, Bleiswijk, The Netherlands).

Illumina sequence libraries of genomic DNA were prepared using the Nextera XT kit (Illumina, San Diego, CA) according to the manufacturer’s instructions, and sequenced on an Illumina MiSeq system with a 500-cycle (2 × 250 bp) MiSeq v2 reagent kit (Illumina). MinION library preparation for barcoded 2D long-read sequencing was performed using the SQK-LSK208 kit (Oxford Nanopore Technologies, Oxford, England, United Kingdom), according to the manufacturer’s instructions, with G-tube (Covaris, Woburn, Massachusetts, United States of America) shearing of chromosomal DNA for 2 x 120 seconds at 1500 *g*. The libraries were sequenced on a MinION sequencer (Oxford Nanopore Technologies) through a SpotON Flow Cell Mk I (R9.4; Oxford Nanopore Technologies).

### Genome assembly and annotation

The quality of the Illumina sequencing data was assessed using FastQC v0.11.5 (https://github.com/s-andrews/FastQC). Illumina sequencing reads were trimmed for quality using nesoni v0.115 (https://github.com/Victorian-Bioinformatics-Consortium/nesoni) using standard settings with the exception of a minimum read length of 100 nucleotides. MinION reads in FastQ format were extracted from Metrichor base-called FAST5-files using Poretools [55]. *De novo* genome hybrid assembly of colistin-susceptible strains was performed using Illumina and Oxford Nanopore data using SPAdes v3.6.2 with the following settings: kmers used: 21, 33, 55, 77, 99, 127, “careful” option turned on and cut-offs for final assemblies: minimum contig/scaffold size of 500bp, and a minimum average scaffold nucleotide coverage of 10 [56,57]. Genome annotation was performed using Prokka [58].

### Phylogenetic analysis, MLST typing, and identification of antibiotic resistance genes

To generate a core genome phylogeny, Illumina/Oxford Nanopore hybrid genome assemblies were aligned using ParSNP v1.2 (37) with 37 publicly available *Klebsiella pneumoniae* complex genomes that cover all phylogroups of the *K. pneumoniae* complex [2]. To include the genome of *K. africanensis* strain 38679, we assembled the genome from raw reads, by processing the raw sequence reads using Nesoni with standard settings, except for minimum read length (75 nucleotides), and subsequent assembly by SPAdes with kmers 21, 33, 55, 77 and the “careful” options turned on.

Figtree was used to visualize and midpoint root the phylogenetic tree (http://tree.bio.ed.ac.uk/). MLST typing was performed using the mlst package v2.10 (https://github.com/tseemann/mlst). Genome assemblies of colistin-susceptible strains were assessed for antibiotic resistance genes by ResFinder 3.1 through standard settings [59].

### Determination of SNPs and indels between axenic colistin-susceptible and colistin-resistant strain pairs

Read-mapping of Nesoni-filtered reads of evolved strains to the genomes of the isogenic colistin-susceptible parental strains was performed using Bowtie2 [60]. SNP and indel-calling was performed using SAMtools 0.1.18 using the following settings: Qscore ≥ 50, mapping quality ≥ 30, a mapping depth ≥ 10 reads, a consensus of ≥ 75% to support a call, and ≥ 1 read in each direction supporting a mutation, as previously described [61]. To correct for potential assembly errors, we also performed the SNP and indel-calling procedure by mapping the reads of the reference isolates against their own assemblies. SNPs and indels found in the reference-versus-reference comparison were ignored in query-versus-reference comparisons. Synonymous mutations were excluded from further analyses. SNPs and indels were manually linked to genes in the assembly.

### Determination of location of IS elements in genomes

To determine which IS elements were present in the genomes of colistin-susceptible strains, we analysed the Illumina/Oxford Nanopore hybrid genome assemblies using ISfinder [62]. Per genome, the IS elements with an E-value < 1e-50 were selected for further study. If multiple distinct IS elements were called at the same position, the element with the highest sequence identity was selected to represent that position.

To detect changes in the position of the identified IS elements, we analysed the genomic assemblies of the isogenic colistin-susceptible and colistin-resistant strain pairs through ISMapper [63]. To maximize the ability of ISMapper to detect IS elements in our sequencing data, the obtained nucleotide sequences of the IS elements in the genome were used as input, and the --cutoff flag of ISMapper was set to 1, whilst other settings remained unchanged. The results were inspected for IS elements that had different positions between the colistin-susceptible, and colistin-resistant strains. Insertion of IS elements was confirmed through PCRs, using DreamTaq Green PCR Master Mix (Thermo Fisher Scientific) and primers spanning the IS insertion site (Supplemental Table S1) and subsequent Sanger sequencing of the PCR product by Macrogen (Amsterdam, The Netherlands).

### SNP and indel calling in evolving populations

To track the genomic changes within the growing cultures under the selective pressure of increasing colistin concentrations, genomic DNA was isolated from the 5-7 overnight cultures of each *in vitro* evolution experiment and sequenced on the Illumina MiSeq platform as described above. SNPs and indels were called as before, with each call supported by at least 25% of reads. Once identified in one or more populations, the abundance of the specific SNPs and indels were then quantified manually for all individual populations of the *in vitro* evolution experiment. Mutations called within 150 bp of a contig end were filtered out, as previously recommended [64]. Identified SNPs and indels were manually linked to genes in the genome assembly, and inspected for synonymous versus non-synonymous mutations. Non-coding mutations were included in subsequent analyses, while synonymous mutations were excluded.

### Determination of growth rate

To determine the maximum specific growth rate, a Bioscreen C instrument (Oy Growth Curves AB, Helsinki, Finland) was used. Overnight cultures were used to inoculate 200 µl fresh LB medium 1:1000. Incubation was set at 37°C with continuous shaking. Growth was observed by measuring the absorbance at 600 nm every 7.5 minutes. Each experiment was performed in triplicate.

### MALDI-TOF analysis of lipid A structures

Isolation of lipid A molecules and subsequent analysis by negative-ion matrix-assisted laser desorption–ionisation time-of-flight (MALDI-TOF) mass spectrometry was performed as previously described [41,65,66]. Briefly, *K. pneumoniae* strains were grown in LB (Oxoid) and the lipid A was purified from stationary cultures using the ammonium hydroxide/isobutyric acid isolation method described earlier [67]. Mass spectrometry analysis were performed on a Bruker autoflex® speed TOF/TOF mass spectrometer in negative reflective mode with delayed extraction using as matrix an equal volume of dihydroxybenzoic acid matrix (Sigma-Aldrich) dissolved in (1:2) acetonitrile-0.1% trifluoroacetic acid. The ion-accelerating voltage was set at 20 kV. Each spectrum was an average of 300 shots. A peptide calibration standard (Bruker) was used to calibrate the MALDI-TOF. Further calibration for lipid A analysis was performed externally using lipid A extracted from *Escherichia coli* strain MG1655 grown in LB medium at 37°C.

### LL-37 survival assay

In order to test the susceptibility of the *K. pneumoniae* strains to LL-37, we adapted previously described protocols [68]. An overnight broth culture was diluted to a concentration of 2.5×10^6^ CFU/ml in 25% LB and incubated with or without the addition of 50 µg/ml LL-37 (AnaSpec Inc, Fermont, California, United States of America) for 90 minutes at 37°C with agitation at 300 rpm in sterile round-bottom 96-well plates (Greiner Bio-One, Alphen aan den Rijn, The Netherlands). After incubation, samples were serially diluted in PBS and plated on LB agar plates. CFUs were counted after overnight incubation at 37°C.

### *Caenorhabditis elegans* virulence assays

*Caenorhabditis elegans* strain CF512 (*rrf-3*(*b26*) *II*; *fem-1*(*hc17*) *IV*), which has a temperature-sensitive reproduction defect, was obtained from the Caenorhabditis Genetics Center at the University of Minnesota, Twin Cities (http://www.cgc.cbs.umn.edu/). CF512 nematodes were maintained at 20°C on Nematode Growth Medium (NGM) agar plates seeded with *E. coli* OP50 [69], and placed on fresh plates at least once per week. For seeding of NGM plates, mid-exponential phase cultures were used. After reaching mid-exponential phase, the cells were washed with PBS, and 1 × 10^6^ CFU were spread on NGM plates, after which the bacterial lawns were grown overnight at 37°C.

To quantify bacterial virulence, *C. elegans* CF512 lifespan assays were performed with synchronized nematodes according to a previously described protocol [70]. For synchronization, nematodes and eggs were collected from a NGM plate in ice-cold filter-sterilized M9 medium, and washed by spinning at 1500 x *g* for 30 seconds [71]. Nematodes were destructed by vigorous vortexing in hypochlorite solution (25 mM NaOH, 1.28% sodium hypochlorite) for two minutes, after which the reaction was stopped by the addition of M9 medium. Eggs were allowed to hatch on NGM plates seeded with *E. coli* OP50 for 6-8 hours at 20°C, after which they were placed at 25°C to avoid progeny. After 48 hours, L3-L4 nematodes were placed on NGM plates (n=40 per plate) seeded with bacterial strains. Plates were scored for live nematodes. Nematodes were considered dead when they did not show spontaneous movement or a response to external stimuli.

### Statistical analysis

Statistical analyses were performed using the parametric one-way ANOVA test with a Dunnett’s test for multiple comparisons (for the determination of maximum growth rates), the non-parametric Mann-Whitney test was used (for the LL-37 survival assay), the parametric one-way ANOVA test with a Sidak’s test for multiple comparisons (for the serum survival assays), and the Mantel-Cox log-rank test (for the *C. elegans* assays). Statistical significance was defined as a p-value < 0.05 for all tests. Statistical analyses were performed using GraphPad Prism 6 software (GraphPad Software, San Diego, California, United States of America).

### Data availability

Sequence data of both the Illumina short-read, and the Oxford Nanopore long-read sequencing has been deposited in the European Nucleotide Archive (accession number PRJEB29521).

## Results

### The colistin-susceptible *K. pneumoniae* complex strains have a diverse genetic background

The four clinical isolates used in this study were obtained from pus, faecal, or urine samples through routine diagnostic procedures in September 2013. All four strains were initially typed as *K. pneumoniae sensu stricto* through routine diagnostic procedures using MALDI-TOF. The susceptibility to colistin of these strains, previously determined in routine diagnostic procedures, was confirmed through antibiotic susceptibility testing using broth microdilution (Figure 1A).

**Figure 1.**
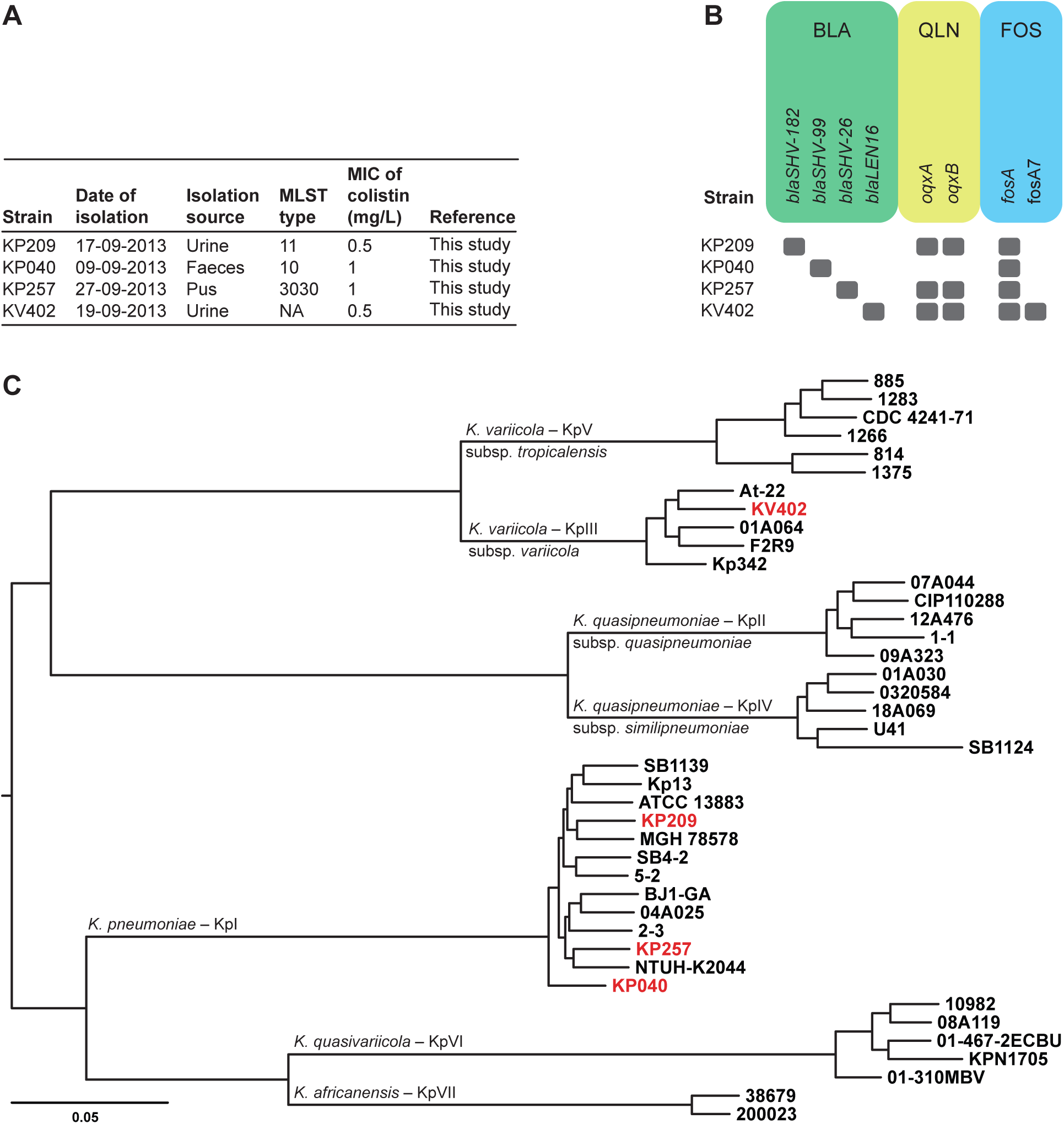
*K. pneumoniae* complex strains: metadata, presence of antibiotic resistance genes, and core-genome phylogenetic analysis. **A)** Overview of the isolates used in this study, including the date and source of isolation, MLST type, and the initial MIC determined. MLST typing of strain KV402 resulted in an incomplete MLST profile, so no conclusive ST could be assigned. NA, not applicable. **B)** Antibiotic resistance genes detected in *K. pneumoniae* complex strains sequenced as part of this study. Classes of antibiotic resistance genes are indicated as follow: BLA, beta-lactam resistance genes; QLN, quinolone resistance genes; FOS, fosfomycin resistance genes. The strains did not carry acquired colistin resistance genes of the *mcr-*family. **C)** Midpoint-rooted phylogenetic tree representing the 1.3-Mbp core-genome alignment of 41 *K. pneumoniae* complex. Taxonomic phylogroups of the *K. pneumoniae* complex [2] are indicated along the branches. The strains used in this study are highlighted in red.

The sequenced genomes of the colistin-susceptible strains were screened for acquired antibiotic resistance genes through ResFinder 3.1 (Figure 1B). None of the nosocomial strains was determined to carry one of the *mcr*-genes. Between two and five acquired antibiotic resistance genes were observed in the genome assemblies, encoding resistance to beta-lactams, quinolones, and fosfomycin.

To accurately identify the phylogenetic position of these nosocomial strains within the *K. pneumoniae* complex, a phylogenetic tree was generated based on the Illumina/Oxford Nanopore hybrid genome assemblies of the colistin-susceptible strains, and 37 publicly available genomes covering all phylogroups in the *K. pneumoniae* complex [2]. Based on a 1.3 Mbp core-genome alignment, the phylogenetic tree showed that strains KP209, KP040, and KP257 clustered in the *K. pneumoniae sensu stricto* (KpI) phylogroup (Figure 1C). Strain KV402 clustered in the *K. variicola* subsp. *variicola* (KpIII) phylogroup, even though it had been typed as *K. pneumoniae sensu stricto* through MALDI-TOF during initial routine diagnostic procedures.

### Colistin resistance emerges through multiple evolutionary trajectories in the *K. pneumoniae complex*

In an effort to understand the evolutionary trajectories through which the *K. pneumoniae* complex strains evolved resistance towards colistin, we deep-sequenced each overnight culture of the *in vitro* evolution experiment (Supplemental Table S2), in which the strains were grown in the presence of increasing concentrations of colistin and identified SNP, indels and excision/integration events of IS elements.

Through these methods, we observed the rapid emergence and fixation of several mutations (Figure 2) in the presence of colistin. In three strains (KP209, KP257, and KV402), these mutations occurred in the genes encoding the PhoPQ two-component regulatory system after one day of culturing (Supplemental Table S3). The PhoPQ two-component regulatory system is a well-known mediator of colistin resistance in *K. pneumoniae* complex strains [30,72]. The G385S substitution identified in PhoQ of KP257 has been previously linked to colistin resistance [73], and the other mutations in *phoPQ* presumably confer colistin resistance to these strains as well. In strain KP040, we observed the integration of an IS*5* element (Supplemental Table S4, Supplemental Data 1) in the promoter region of both the *ccrAB* operon, which encodes a two-component regulatory system, that has previously been linked to colistin resistance [31], and the CrrAB-controlled *crrC* gene, which encodes an activator of the PmrAB two-component regulatory system [74]. In addition, an intergenic SNP (located in promoter regions of *ecpR* or *phnC*) in strain KP040 became fixed in the population on the first day of culturing. Both EcpR and PhnC have not previously been associated with colistin resistance. Although other mutations, in other locations, also occurred during the first day of culturing, these mutations failed to become fixed in the population, and were either lost on subsequent days, or did not change in abundance over time.

**Figure 2.**
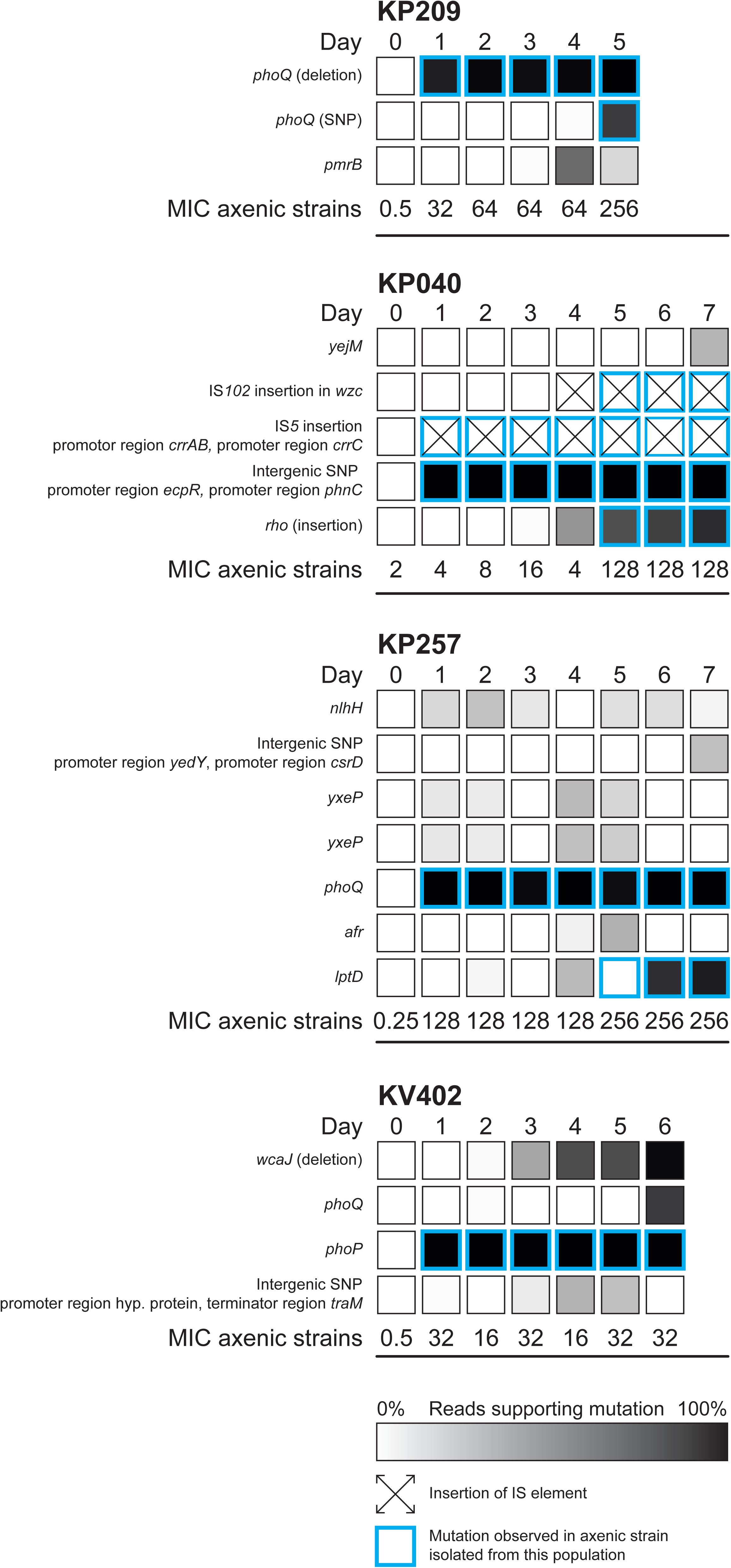
Population analysis of mutations during *in vitro* evolution in the presence of colistin. For each strain, and each day of the *in vitro* evolution experiment, the positions that have mutated compared to the colistin-susceptible strain are indicated. For SNPs and indels, the number of reads supporting a mutation at a given location was used to estimate the abundance of the mutation. Novel integrations of IS elements are also indicated. For mutations not located in a coding sequence, nearby coding sequences are indicated. Mutations and IS element integrations observed in the axenic, strain isolated daily from each population are indicated by a blue border. The MIC of colistin for each axenic strain isolated from the *in vitro* evolution population is indicated. The MIC values represent the mode from three independent experiments performed in duplo. Hyp. protein: hypothetical protein.

On subsequent days of the *in vitro* evolution experiment, novel mutations in the populations were associated with additional increases in MIC of colistin. New SNPs that were fixed in the populations were observed in *phoQ* (strain KP209 (day 5), and KV402 (day 6)), and *pmrB* (KP209 (day4)). In strain KP257, a SNP in *lptD* was first observed on day 3, and was then fixed in the population. The *lptD* gene encodes a barrel-shaped transporter that transports LPS onto the outer leaflet of the outer membrane. Mutations in genes located in the K-locus, involved in capsule synthesis, were also detected. In strain KV402 a 13-bp deletion was observed in *wcaJ* from day 3 onwards, leading to a premature stop-codon. In KP040 a new insertion of IS*102*, inactivating *wzc* was observed from day 4. In addition, a 12-bp insertion in the gene encoding the Rho transcription termination factor was observed in KP040. We did not observe any mutations in the *mgrB* gene, encoding the negative regulator of the PhoPQ two-component regulatory system, in these *in vitro* evolution experiments.

### *K. pneumoniae* can rapidly develop colistin resistance without loss of fitness

To characterize changes in fitness and virulence characteristics as a result of development of colistin resistance, we isolated an axenic strain on each day of the *in vitro* evolution experiments. The genome sequences of the axenic strains of the last day of the *in vitro* evolution experiments were determined by Illumina sequencing. SNPs, indels and IS element insertions were identified in these strains in comparison with the colistin-susceptible parental strain. After combining these data with the population sequencing data described above, we determined the presence of these mutations in the axenic strains isolated after each day of the *in vitro* evolution experiment by targeted PCRs and Sanger sequencing of the amplicons. We were thus able to correlate the occurrence of mutations with increases in the MIC of colistin in each strain.

All four strains developed levels of resistance to colistin above the breakpoint value after one overnight incubation of the colistin-susceptible strain in the presence of the antibiotic (Figure 2). The initial mutations in *phoPQ* were associated with an increase in MIC in strains KP209, KP257, and KV402 (Figure 2). The integration of the IS*5* element in the promoter region of *crrAB* and *crrC*, and the appearance of an intergenic SNP between *ecpR* and *phnC*, also occur simultaneously with an increase in the MIC of colistin. The additional SNP in *phoQ* in KP209 was not associated with an increase in the MIC of colistin. Integration of IS*102* in *wzc* of the K-locus, as well as the 12-bp insertion in the gene encoding the transcription termination factor Rho, was associated with an additional increase in the MIC of colistin in strain KP040. The SNP in *lptD* in strain KP257 did not lead to a meaningful increase in the MIC of colistin. The culture isolated from the last day of the KV402 *in vitro* evolution experiments had a SNP in *yciM* (Supplemental Table S5), encoding a negative regulator of LPS biosynthesis, but this did not contribute to a further reduced susceptibility to colistin.

The measurement of the maximum growth rate as a proxy for general fitness of the axenic strains isolated on the different days of the *in vitro* evolution experiment showed that the increase in MIC of colistin to values above 2 µg/ml after one overnight incubation, did not negatively affect the maximum growth rate for strains KP209, KP040, and KV402. Only the initial increase in MIC of colistin in strain KP257 had a negative impact on the maximum growth rate, decreasing the maximum growth rate by 37% (Figure 3). Over time, the maximum growth rates of strains KP209 and KV402 decreased 13.4% and 9.5%, respectively, compared to the maximum growth rate of the colistin-susceptible strain. In strain KP040, an increase of 10.0% in maximum growth rate was observed during the course of the *in vitro* evolution experiment.

**Figure 3.**
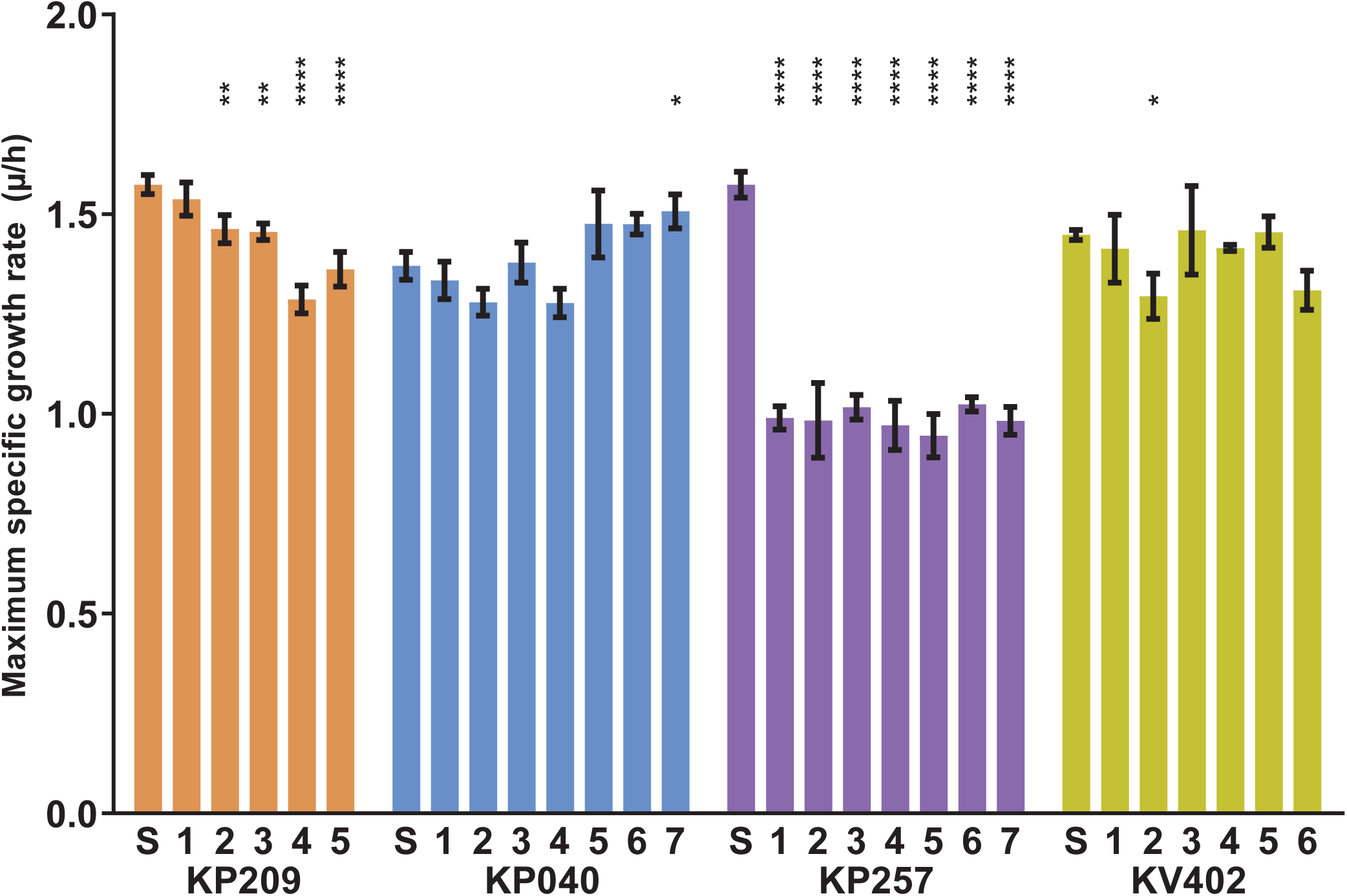
Maximum growth rate of colistin resistant evolved strains. Optical density at 600 nm (OD_600_) was measured every 7.5 minutes. Representative data of three individual experiments, performed in triplicate are shown. Mean and standard deviations are shown. A parametric one-way ANOVA with Dunnett’s multiple correction was used for the statistical analysis of the differences in growth rates between the axenic strains isolated from each day of the *in vitro* evolution experiment and the colistin-susceptible parental strain. Outcomes of the statistical analysis are indicated by asterisks: p < 0.05 (*), < 0.01 (**), <0.001 (***), or <0.0001 (****).

### Colistin-resistant *K. pneumoniae* complex strains have lipid A that is modified through hydroxylation, palmitoylation and addition of 4-amino-4-deoxy-L-arabinose (L-Ara4N)

To determine the modifications to lipid A in the colistin-resistant strains, we performed MALDI-TOF analysis on lipid A isolated from the colistin-susceptible strain, and the axenic strain of the last day of the *in vitro* evolution experiments. The MALDI-TOF spectra of lipid A isolated from colistin-susceptible strains (Figure 4A), showed a dominant peak from hexa-acylated lipid A (mass-to-charge ratio (*m/z*) 1824), corresponding to two glucosamines, two phosphates, four 3-OH-C_14_ and two C_14_ acyl chains [65]. Additional minor peaks in the MALDI-TOF spectrum of the susceptible strains could be observed at *m/z* 1840, corresponding to the hydroxylation (*m/z* 16) of one of the C_14_ acyl-groups of hexa-acylated lipid A (*m/z* 1824), and at *m/z* 2063 (in KP209 and KP257), corresponding to a hepta-acylated lipid A, with an additional acylation of lipid A (*m/z* 1824) with a palmitoyl group (*m/z* 239).

**Figure 4.**
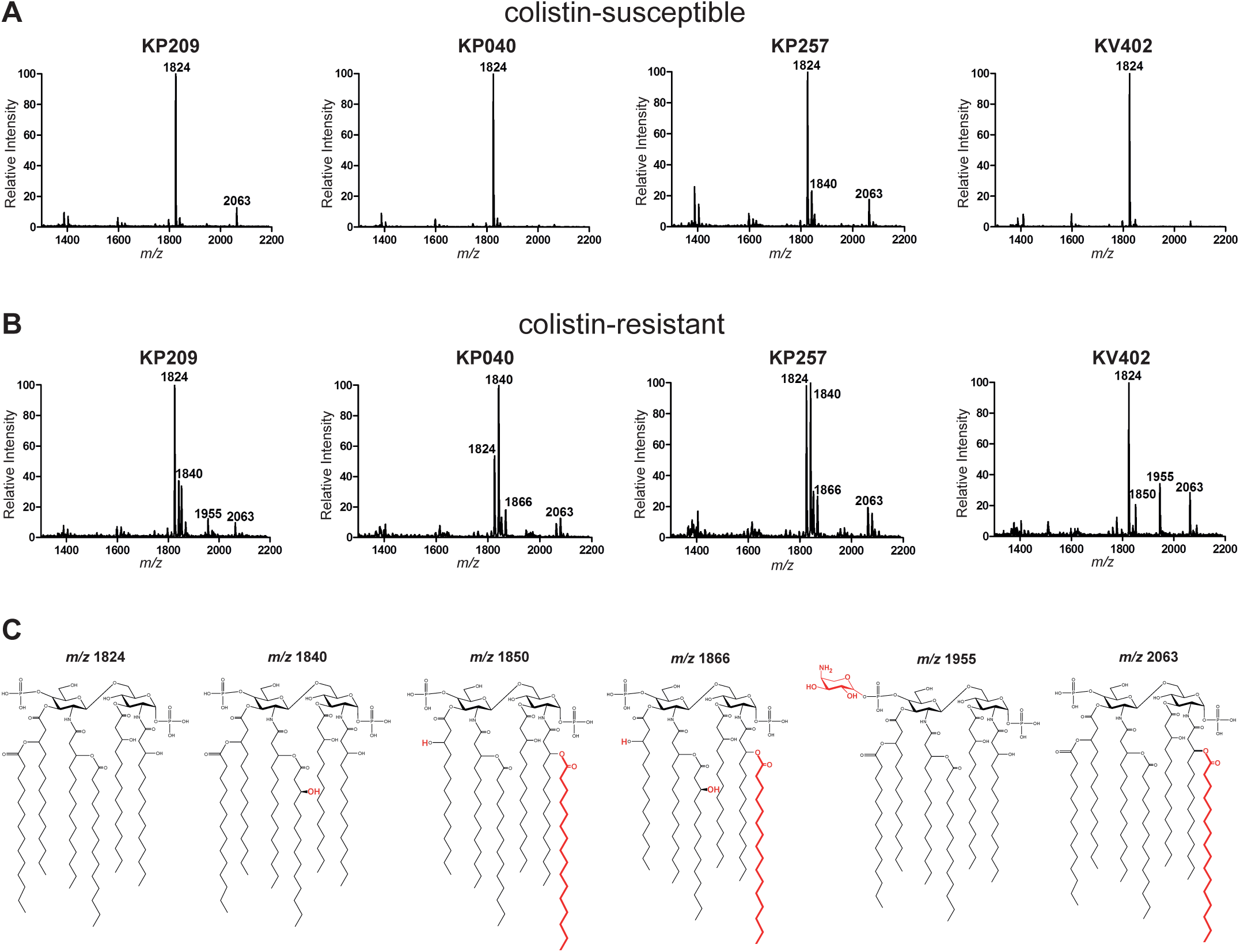
Lipid A modifications in colistin-susceptible and colistin-resistant strains. MALDI-TOF spectra showing the mass-to-charge (m/z) ratio values of the isolated lipid A from (A) colistin-susceptible, and (B) colistin-resistant axenic strains, isolated from the cultures of the last day of the *in vitro* evolution experiment. C) Proposed chemical structures of lipid A-moieties corresponding to the observed m/z-values in the MALDI-TOF spectra. Modifications relative to the unmodified hexa-acylated lipid A corresponding to m/z value 1824 are depicted in red. Hydroxylation of an acyl-chain adds 16 to the m/z ratio, 4-amino-4-deoxy-L-arabinose adds 131, acylation with palmitate adds 239.

All the MALDI-TOF spectra of lipid A isolated from colistin-resistant strains show additional peaks (Figure 4B), indicating the modification of their lipid A. In the spectra of colistin-resistant KP209 and KV402, lipid A *m/z* 1955 was observed, indicating addition of L-Ara4N (*m/z* 131) to the hexa-acylated lipid A *m/z* 1824. In colistin-resistant KV402 lipid A *m/z* 1850 was observed, consistent with hexa-acylated lipid A *m/z* 1824 with one C_16_ acyl chain (Figure 4C). The peak at *m/z* 1866 in the MALDI-TOF spectra of colistin-resistant KP040 and KP257 was consistent with hydroxylation of lipid A *m/z* 1850.

### Development of colistin resistance is associated with increased LL-37 resistance and virulence in a *C. elegans* survival model

Since the mechanisms of LPS modification that result in colistin resistance and immune evasion are similar [75], we investigated the effects of colistin resistance on virulence characteristics. Since LL-37 is a human cathelicidin antimicrobial peptide, it has a similar mechanism of action as colistin. Previously, cross-resistance between colistin and LL-37 was observed in *Acinetobacter baumannii* [68]. To assess the possible cross-resistance between colistin and LL-37, we determined the survival of the colistin-resistant strains under influence of LL-37. We observed that three of the four colistin-resistant strains (KP209, KP040, and KP257) showed a decreased susceptibility to killing by LL-37 compared to their colistin-susceptible parental strains (Figure 5). In contrast, development of colistin resistance in strain KV402 did not affect susceptibility to LL-37.

**Figure 5.**
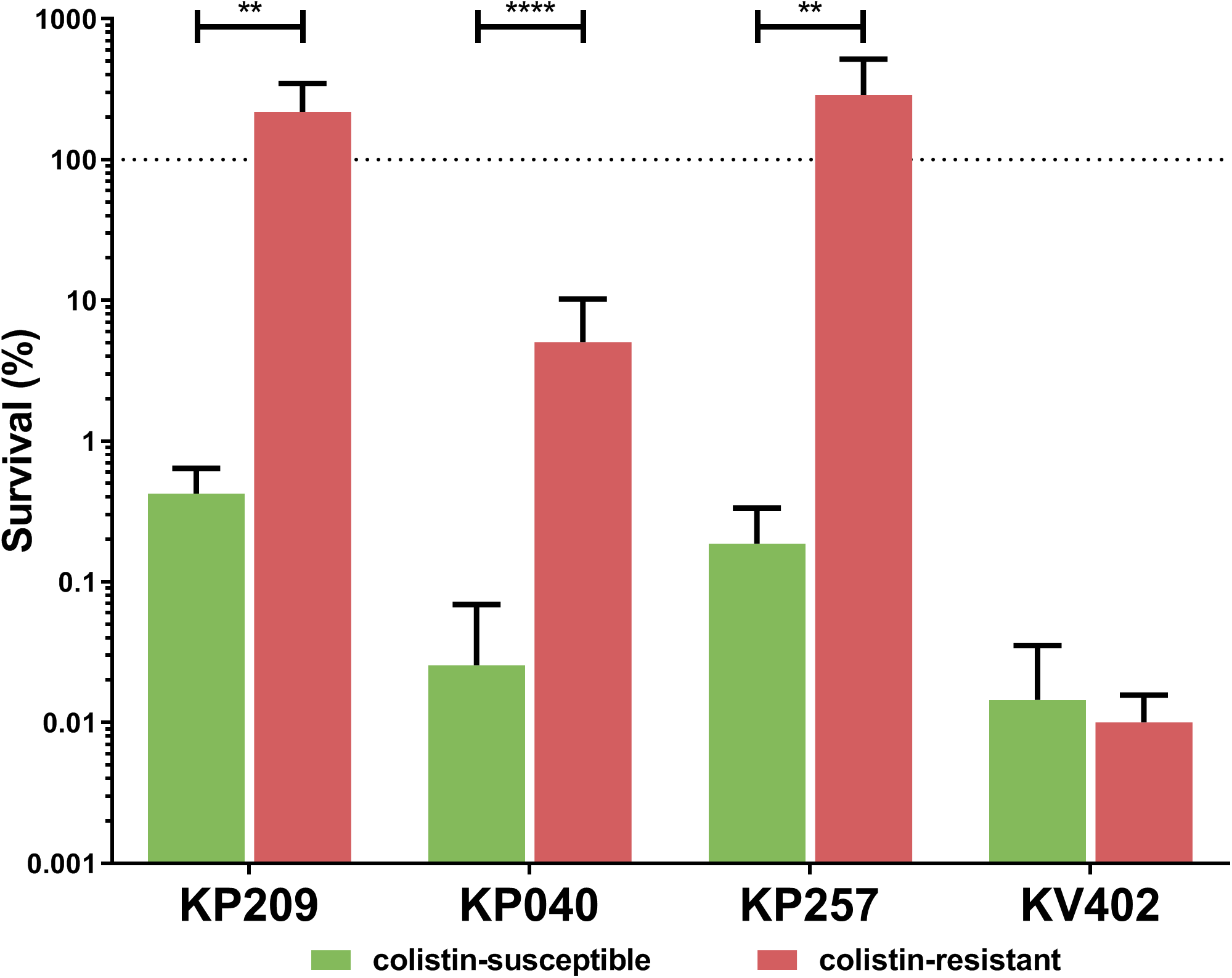
Susceptibility of colistin-susceptible and colistin-resistant strains to the human cathelicidin LL-37. Strains were incubated for 90 minutes in 25% LB at 37ºC with or without the addition of 50 µg/ml LL-37. Viability was assessed by determination of the number of colony-forming units. The non-parametric Mann-Whitney test was used as statistical test and significance was defined as a p-value of < 0.05 (*), < 0.01 (**), <0.001 (***), or <0.0001 (****).

To investigate the possible consequences of colistin resistance on virulence, we exposed the nematode *C. elegans* strain CF512 to the colistin-susceptible/resistant strain pairs. *C. elegans* had a decreased lifespan on a lawn of colistin-resistant KP209 (Figure 6) and KP040, compared to their colistin-susceptible strains. Survival of *C. elegans* was not affected by growth on colistin-resistant strains derived from KP257 and KV402, compared to the colistin-susceptible parental strains.

**Figure 6.**
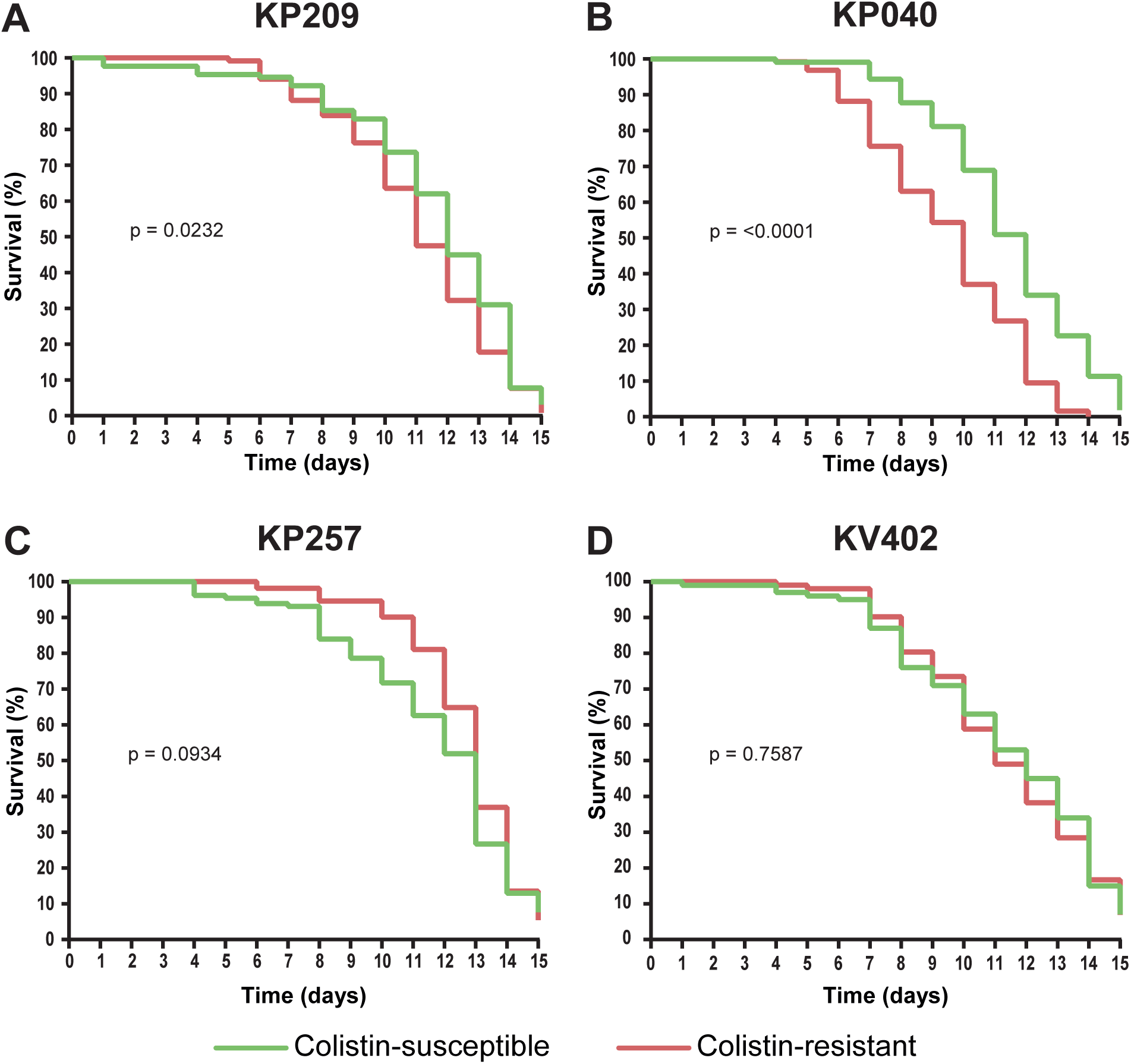
Survival of *C. elegans* on lawns of *K. pneumoniae* colistin-susceptible and colistin-resistant strains complex strains. *C. elegans* CF512 were kept on a lawn of colistin-susceptible (green) and colistin-resistant (red) *K. pneumoniae* complex strains. Survival was scored over a period of 15 days. The data represent three independent experiments in which a total of 129 (in colistin-susceptible KP209), 118 (colistin-resistant KP209), 106 (colistin-suscptible KP040), 127 (colistin-resistant KP040), 127 (colistin-susceptible KP257), 131 (colistin-resistant KP257), 100 (colistin-susceptible KP402), and 102 (colistin-resistant KP402) *C. elegans* nematodes were used. Statistical significance according to Mantel-Cox log-rank test is indicated. Statistical significance was defined as a p-value < 0.05.

## Discussion

Colistin plays a pivotal role in public health due to its last-resort status for treatment of infections with multidrug-resistant Gram-negative bacteria. The increasing number of reports of *K. pneumoniae* strains that have acquired resistance to multiple antibiotics, including colistin, is thus a cause for increasing concern [7–9,29]. In this study, we observed the swift development of colistin resistance through diverse evolutionary trajectories by conducting an *in vitro* evolution experiment with nosocomial *K. pneumoniae* complex strains. Development of colistin resistance had no, or only a minor, impact on maximum growth rate in three out of four *in vitro* evolution experiments performed here. This suggests that colistin may rapidly lose its effectiveness in the treatment of infections caused by multidrug-resistant *K. pneumoniae* complex strains as fitness costs associated with colistin resistance seem limited.

We observe that mutations associated with an increase in MIC of colistin seem confined to genes from functional groups involved in the synthesis and modification of LPS, and the synthesis of capsular polysaccharides, which are both important surface-associated structures. In the genes encoding the PhoPQ two-component regulatory system, which have a role in regulating modifications of LPS and contribute to colistin resistance in Enterobacteriaceae [30,34,76], we found variations in both PhoP (a D191N substitution), and PhoQ (a G385S substitution, and a 12-bp deletion). The G385S PhoQ substitution has previously been described in a clinical strain of colistin-resistant *K. pneumoniae* [73]. Outside PhoPQ, we found that a novel integration of an IS*5* element in the promoter region associated with the genes encoding CrrAB and CcrC coincides with increase in MIC of colistin. The IS*5* element can influence the transcriptional activity of the genes located near its integration site [77]. The activity of PmrAB may be influenced by CrrAB through CrrC [31,74,78]. In line with previous observations, in which insertions of IS elements were associated with resistance to tigecycline and colistin, we hypothesize that the insertion of IS5 may lead to increased expression of either CrrAB and/or CrrC, and cause colistin resistance [40,79].

For the genes involved in capsule synthesis, we observe that the inactivation of *wzc* of the K-locus, by the IS*102* element coincides with an increase in the MIC of colistin. In *E. coli*, Wzc is involved in the synthesis and export of extracellular polysaccharides containing colanic acid [80], but also the phosphorylation of other endogenous proteins [81]. Wzc has previously been hypothesised to be involved in colistin resistance in *E. coli*, and it may act similarly in *K. pneumoniae* [81–83]. The loss of Wzc may potentially cause colistin resistance through two mechanisms. A reduction in the export of colanic acid units (the building blocks of *K. pneumoniae* capsule), can lead to the accumulation of colanic acid metabolic intermediates, including UDP-glucuronic acid. This accumulation has been hypothesised to lead to an increased flux towards biosynthesis of UDP-L-Ara4N, resulting in the modification of lipid A with L-Ara4N [84]. Alternatively, the absence or reduction of negatively charged colanic acid residues on the cell surface could lower local concentrations of positively charged colistin molecules, thereby reducing damage to the outer membrane [84]. Together with the inactivation of *wzc*, we observe a 12-bp insertion in the highly-conserved *rho* gene, encoding the transcription termination factor Rho. Rho has not been previously linked to colistin resistance, but mutations in *rho* may have pleiotropic effects on transcription [85], which could influence the expression of genes involved in, or may compensate for fitness costs caused by, colistin resistance.

Notably, we did not find any alterations in *mgrB*, which is an otherwise important mechanism through which colistin resistance may occur in nosocomial *K. pneumoniae* complex strains [25,36–39]. Nevertheless colistin-resistant clinical *K. pneumoniae* isolates without mutations in *mgrB* are also frequently encountered [73,74,86–89]. We can only speculate on the reasons for the absence of *mgrB* mutations in our *in vitro* evolution experiments, but the relatively short duration of this experiment performed with a limited number of strains, likely implies that we have not covered all potential colistin resistance mechanisms in *K. pneumoniae*.

In this study, we observe the development and fixation of several mutations in systems that are known to contribute to colistin resistance. We could not however, generate targeted mutants within the clinical strains. The multidrug-resistant nature of these clinical strains, prevents the use sof often-used systems to generate mutations. The impact of developing colistin resistance through the observed mutations might extend past the inability to treat the infection through antibiotic therapy, as modifications to lipid A may reduce the susceptibility to antimicrobial peptides, or increase virulence, as we show in this study [41,65,68,90]. However, the mechanisms behind the differential effects on virulence of colistin resistance in the *K. pneumoniae* complex are currently not fully understood and are deserving of further study. A single *K. variicola* isolate was included in this study. While *K. variicola* can cause life-threatening infections in immunocompromised individuals [5], it remains currently understudied. Additional studies into the mechanisms of colistin resistance and their impact on fitness and virulence may be warranted in this species.

To prevent the rapid emergence of colistin resistance in *K. pneumoniae* complex strains in clinical settings, the use of colistin in synergistic combinations with other antibiotics may limit development of resistance [91]. Additionally, colistin resistance in *K. pneumoniae* may confer collateral sensitivity to other classes of antibiotics, and may yield combinations of antibiotics that can be used alternately, in a process termed drug cycling [92,93]. More in-depth knowledge about colistin resistance mechanisms may also facilitate the development of novel therapeutic targets. Although a diverse set of two-component systems may be implicated in the development of resistance, the PmrAB two-component system appears to play a central role since both the PhoPQ and CrrAB two-component systems activate PmrAB, through PmrD and CrrC respectively [74]. The development of a small-molecule inhibitor targeting the two-component systems involved in lipid A modifications [94], may be essential to lengthen the clinical lifespan of colistin as a last-resort drug in treatment of *K. pneumoniae* complex strains.

## Acknowledgements

We thank the Utrecht Sequence Facility and Ivo Renkens for their expertise in MinION Oxford Nanopore sequencing, Lidewij W. Rümke for the contribution of clinical metadata of the used nosocomial isolates, Dr. Inge The for advice on *C. elegans* assays, and Dr. Evelien T.M. Berends for helpful discussions.

## Author contributions

A.B.J., D.J.D. and G.M. performed experiments and analysed data. A.B.J. and M.R.C.R performed bioinformatic analyses. S.H.M.R., J.A.B., and W.v.S designed experiments. A.B.J., M.J.M.B, S.H.M.R., R.J.L.W., J.A.B. and W.v.S. wrote the manuscript. All authors reviewed and approved the manuscript prior to submission.

---

**Supplemental Table S1**

**Sequences of oligonucleotide primers.** The name indicates the specific strain and target site of the primer. The nucleic acid sequence (5’-3’) is indicated.

**Supplemental Table S2**

**Summary of the read data from the sequencing runs, and genome assembly information.** Per strain, the sequenced samples are indicated. The axenic colistin-resistant strain was picked from the last day of the *in vitro* evolution experiment. For the colistin-susceptible parental strain, the assembly statistics for the Illumina/Oxford Nanopore hybrid genome assembly are indicated. For all samples, the number of reads mapped to the hybrid assembly of the colistin-susceptible parental strain, and the average coverage are indicated. NA, not applicable

**Supplemental Table S3**

**Mutations observed in the populations of the *in vitro* evolution experiment.** The positions that were observed to have mutated through a SNP or indel in the populations cultured in the presence of colistin, compared to the colistin-susceptible strain, are indicated. For each SNP or indel, the strain in which it was found, the associated genes, the location in the specific scaffold of the Illumina/Oxford Nanopore hybrid genome assembly, the reference allele, and the mutant allele are indicated. If the mutation was observed in a coding sequence, the change in the coding sequence of the affected gene is indicated. The residues are numbered according to their position in the reference allele. For mutations not located in a coding sequence, the associated genes are indicated. Hyp. protein; hypothetical protein.

**Supplemental Table S4**

**Overview of location of IS elements.** Position of IS elements as determined by ISMapper in the colistin-susceptible and colistin-resistant strains (isolated on the last day of the *in vitro* evolution experiments). The result of ISMapper was classified as excision, novel integration, or stable per IS element. Events were classified compared to the Illumina/Oxford Nanopore hybrid genome assembly of the colistin-susceptible strain. For IS elements with an excision, or novel integration, the changed position was checked through targeted PCRs (Supplemental Figure S1) and Sanger sequencing in all isolates collected during the *in vitro* evolution experiment. NA: not applicable.

**Supplemental Table S5**

**Mutations observed in whole-genome sequence of axenic colistin-resistant strains.** For each strain, the positions that have been mutated between the colistin-susceptible strain, and the colistin-resistant strain picked from the last day of culturing are indicated. For each mutation, the associated feature, the location in the specific scaffold of the Illumina/Oxford Nanopore hybrid genome assembly are indicated, the reference allele and the mutant allele are indicated. If the mutation was observed in a coding sequence, the changes in the coding sequence of the affected gene are indicated. The residues are numbered according to their position in the reference allele. For mutations not located in a coding sequence, the associated coding sequences are indicated.

**Supplemental Figure S1**

**Design of, and concentration of colistin used during, the *in vitro* evolution experiment.**

*In vitro* evolution to colistin resistance was achieved by culturing in increasing colistin concentrations over a period of 5-7 days. Prior to the *in vitro* evolution experiments, MICs to colistin were determined in LB. Each strain was grown in 1 ml LB with initial colistin concentrations of 1 and 2 times the MIC. After overnight growth, 1 μl of the cultures with the highest concentration of colistin that had visible growth were used to propagate a fresh culture by inoculating 1 ml of fresh LB, supplemented with the same or twice the concentration of colistin in which growth was observed in the previous day’s culture. This process was repeated for 5-7 days. Each overnight culture was stored at −80°C in 20% glycerol. B) Colistin concentrations used during *in vitro* evolution experiments. A green background indicates the observation of growth after overnight incubation at 37ºC, red indicates no observable growth. The cultures from which the concentration is underlined, were used to propagate the *in vitro* evolution experiment. If no growth was observed in both cultures, cultures were re-inoculated from −80°C stocks of the susceptible strain (only when no growth was observed on day 1), or from cultures stored from the previous day.

**Supplemental Figure S2**

**Agarose gel electrophoresis to determine integration sites of IS elements**. PCR reactions were performed to validate the ISMapper results suggesting IS element excision or integration between the colistin-susceptible parental strains and the colistin-resistant strains of strains KP209 (**A**), KP040 (**B**), and KP257 (**C**), from the last day of the *in vitro* evolution experiment. PCR reactions were performed using primers spanning the integration site of each IS element (Supplemental Table S1). The 1 kb plus DNA ladder (Thermo Scientific) was used for size comparison. The order of samples in each agarose gel electrophoreses is: 1 kb plus DNA ladder, colistin-susceptible strain, axenic strain of the last day of the in vitro evolution experiment, 1 kb plus ladder, colistin susceptible strain, axenic strain for each day of the in vitro evolution experiment, 1 kb plus ladder, negative water control. The order for IS*Ehe3* was: 1 kb plus marker, colistin susceptible strain, axenic strain for each day of the in vitro evolution experiment, 1 kb plus ladder colistin susceptible strain, axenic strain for each day of the *in vitro* evolution experiment, 1 kb plus ladder.

## References

1. Holt KE, Wertheim H, Zadoks RN, Baker S, Whitehouse CA, Dance D, et al. Genomic analysis of diversity, population structure, virulence, and antimicrobial resistance in *Klebsiella pneumoniae*, an urgent threat to public health. Proc Natl Acad Sci U S A. 2015;112(27):E3574–81.

2. Rodrigues C, Passet V, Rakotondrasoa A, Diallo TA, Criscuolo A, Brisse S. Description of *Klebsiella africanensis* sp. nov., *Klebsiella variicola* subsp. *tropicalensis* subsp. nov. and *Klebsiella variicola* subsp. *variicola* subsp. nov. Res Microbiol. 2019;170(3):165–70.

3. Maatallah M, Vading M, Humaun Kabir M, Bakhrouf A, Kalin M, Nauclér P, et al. *Klebsiella variicola* is a frequent cause of bloodstream infection in the Stockholm area, and associated with higher mortality compared to *K. pneumoniae*. PLoS One. 2014;9(11):e113539.

4. Mathers AJ, Crook D, Vaughan A, Barry KE, Vegesana K, Stoesser N, et al. *Klebsiella quasipneumoniae* provides a window into carbapenemase gene transfer, plasmid rearrangements, and patient interactions with the hospital environment. Antimicrob Agents Chemother. 2019;63(6):1–12.

5. Rodríguez-Medina N, Barrios-Camacho H, Duran-Bedolla J, Garza-Ramos U. *Klebsiella variicola*: an emerging pathogen in humans. Emerg Microbes Infect. 2019;8(1):973–88.

6. World Health Organization. Antimicrobial resistance. Global report on surveillance. 2014. European Centre for Disease Prevention and Control. Surveillance of antimicrobial resistance in Europe 2018. Surveillance of antimicrobial resistance in Europe. 2019.

7. Monaco M, Giani T, Raffone M, Arena F, Garcia-Fernandez A, Pollini S, et al. Colistin resistance superimposed to endemic carbapenem-resistant *Klebsiella pneumoniae*: a rapidly evolving problem in Italy, November 2013 to April 2014. Eurosurveillance. 2014;19(42):20939.

8. Parisi SG, Bartolini A, Santacatterina E, Castellani E, Ghirardo R, Berto A, et al. Prevalence of *Klebsiella pneumoniae* strains producing carbapenemases and increase of resistance to colistin in an Italian teaching hospital from January 2012 to December 2014. BMC Infect Dis. 2015;15:244.

9. World Health Organization. 2019 Antibacterial agents in clinical development: an analysis of the antibacterial clinical development pipeline. 2019.

10. Shore CK, Coukell A. Roadmap for antibiotic discovery. Nat Microbiol. 2016;1(6):16083.

11. Llaca-Díaz JM, Mendoza-Olazarán S, Camacho-Ortiz A, Flores S, Garza-González E. One-year surveillance of ESKAPE pathogens in an intensive care unit of Monterrey, Mexico. Chemotherapy. 2013;58(6):475–81.

12. Klein EY, Van Boeckel TP, Martinez EM, Pant S, Gandra S, Levin SA, et al. Global increase and geographic convergence in antibiotic consumption between 2000 and 2015. Proc Natl Acad Sci. 2018;115(15):E3463–70.

13. Ozkan G, Ulusoy S, Orem A, Alkanat M, Mungan S, Yulug E, et al. How does colistin-induced nephropathy develop and can it be treated? Antimicrob Agents Chemother. 2013;57(8):3463–9.

14. Falagas ME, Kasiakou SK. Colistin: the revival of polymyxins for the management of multidrug-resistant Gram-negative bacterial infections. Clin Infect Dis. 2005 May 1;40(9):1333–41.

15. Lim LM, Ly N, Anderson D, Yang JC, Macander L, Jarkowski A, et al. Resurgence of colistin: a review of resistance, toxicity, pharmacodynamics, and dosing. Pharmacotherapy. 2010 Dec;30(12):1279–91.

16. Velkov T, Thompson PE, Nation RL, Li J. Structure-activity relationships of polymyxin antibiotics. J Med Chem. 2010;53(5):1898–916.

17. Domingues MM, Inácio RG, Raimundo JM, Martins M, Castanho MARB, Santos NC. Biophysical characterization of polymyxin B interaction with LPS aggregates and membrane model systems. Biopolymers. 2012 Jul 16;98(4):338–44.

18. Landman D, Georgescu C, Martin DA, Quale J. Polymyxins revisited. Clin Microbiol Rev. 2008 Jul;21(3):449–65.

19. Blair JMA, Webber MA, Baylay AJ, Ogbolu DO, Piddock LJV. Molecular mechanisms of antibiotic resistance. Chem Commun. 2011;47(14):4055–61.

20. Putker F, Bos MP, Tommassen J. Transport of lipopolysaccharide to the Gram-negative bacterial cell surface. FEMS Microbiol Rev. 2015;39(6):985–1002.

21. Sabnis A, Klöckner A, Becce M, Evans LE, Furniss RCD, Mavridou DAI, et al. Colistin kills bacteria by targeting lipopolysaccharide in the cytoplasmic membrane. bioRxiv. 2018;479618.

22. Walsh TR, Wu Y. China bans colistin as a feed additive for animals. Lancet Infect Dis. 2016;16(10):1102–3.

23. Timmerman T, Dewulf J, Catry B, Feyen B, Opsomer G, Kruif A de, et al. Quantification and evaluation of antimicrobial drug use in group treatments for fattening pigs in Belgium. Prev Vet Med. 2006;74(4):251–63.

24. Halaby T, Kucukkose E, Janssen AB, Rogers MRC, Doorduijn DJ, van der Zanden AGM, et al. Genomic characterization of colistin heteroresistance in *Klebsiella pneumoniae* during a nosocomial outbreak. Antimicrob Agents Chemother. 2016;60(11):6837–43.

25. Kieffer N, Aires-de-Sousa M, Nordmann P, Poirel L. High rate of MCR-1–producing *Escherichia coli* and *Klebsiella pneumoniae* among pigs, Portugal. Emerg Infect Dis. 2017;23(12):2023–9.

26. Wang X, Liu Y, Qi X, Wang R, Jin L, Zhao M, et al. Molecular epidemiology of colistin-resistant Enterobacteriaceae in inpatient and avian isolates from China: high prevalence of *mcr*-negative *Klebsiella pneumoniae*. Int J Antimicrob Agents. 2017;50(4):536–41.

27. Tuo H, Yang Y, Tao X, Liu D, Li Y, Xie X, et al. The prevalence of colistin resistant strains and antibiotic resistance gene profiles in Funan river, China. Front Microbiol. 2018;9:3094.

28. Elemam A, Rahimian J, Mandell W. Infection with panresistant *Klebsiella pneumoniae*: a report of 2 cases and a brief review of the literature. Clin Infect Dis. 2009;49(2):271–4.

29. Olaitan AO, Morand S, Rolain J-M. Mechanisms of polymyxin resistance: acquired and intrinsic resistance in bacteria. Front Microbiol. 2014 Nov 26;5:643.

30. Wright MS, Suzuki Y, Jones MB, Marshall SH, Rudin SD, van Duin D, et al. Genomic and transcriptomic analyses of colistin-resistant clinical isolates of *Klebsiella pneumoniae* reveal multiple pathways of resistance. Antimicrob Agents Chemother. 2015;59(1):536–43.

31. Liu YY, Wang Y, Walsh TR, Yi LX, Zhang R, Spencer J, et al. Emergence of plasmid-mediated colistin resistance mechanism MCR-1 in animals and human beings in China: a microbiological and molecular biological study. Lancet Infect Dis. 2016;16(2):161–8.

32. Nang SC, Li J, Velkov T. The rise and spread of mcr plasmid-mediated polymyxin resistance. Crit Rev Microbiol. 2019;45(2):131–61.

33. Poirel L, Jayol A, Nordmann P. Polymyxins: antibacterial activity, susceptibility testing, and resistance mechanisms encoded by plasmids or chromosomes. Clin Microbiol Rev. 2017;30(2):557–96.

34. Sun J, Zhang H, Liu YH, Feng Y. Towards understanding MCR-like colistin resistance. Trends Microbiol. 2018;26(9):794–808.

35. Cannatelli A, Giani T, D’Andrea MM, Di Pilato V, Arena F, Conte V, et al. MgrB inactivation is a common mechanism of colistin resistance in KPC carbapenemase-producing *Klebsiella pneumoniae* of clinical origin. Antimicrob Agents Chemother. 2014 Jul 14;58(July):5696–5703.

36. Jayol A, Poirel L, Villegas M-V, Nordmann P. Modulation of *mgrB* gene expression as a source of colistin resistance in *Klebsiella oxytoca*. Int J Antimicrob Agents. 2015 Mar 28;46(1):108–10.

37. Cannatelli A, Santos-Lopez A, Giani T, Gonzalez-Zorn B, Rossolini GM. Polymyxin resistance caused by *mgrB* inactivation is not associated with significant biological cost in *Klebsiella pneumoniae*. Antimicrob Agents Chemother. 2015 Feb 17;59(5):2898–900.

38. Aires CAM, Pereira PS, Asensi MD, Carvalho-Assef APD. MgrB mutations mediating polymyxin B resistance in *Klebsiella pneumoniae* isolates from rectal surveillance swabs in Brazil. Antimicrob Agents Chemother. 2016;60(11):6969–72.

39. Yang T, Wang S, Lin J-E, Griffith BTS, Lian S, Hong Z, et al. Contributions of insertion sequences conferring colistin resistance in *Klebsiella pneumoniae*. Int J Antimicrob Agents. 2020;53(6):105894.

40. Kidd TJ, Mills G, Sá-Pessoa J, Dumigan A, Frank CG, Insua JL, et al. A *Klebsiella pneumoniae* antibiotic resistance mechanism that subdues host defences and promotes virulence. EMBO Mol Med. 2017;9(4):430–47.

41. Ni W, Li Y, Guan J, Zhao J, Cui J, Wang R, et al. Effects of efflux pump inhibitors on colistin resistance in multidrug-resistant Gram-negative bacteria. Antimicrob Agents Chemother. 2016;60(5):3215–8.

42. Padilla E, Llobet E, Doménech-Sánchez A, Martínez-Martínez L, Bengoechea JA, Albertí S. *Klebsiella pneumoniae* AcrAB efflux pump contributes to antimicrobial resistance and virulence. Antimicrob Agents Chemother. 2010 Jan;54(1):177–83.

43. Mahalakshmi S, Sunayana MR, Saisree L, Reddy M. *yciM* is an essential gene required for regulation of lipopolysaccharide synthesis in *Escherichia coli*. Mol Microbiol. 2014;91(1):145–57.

44. Llobet E, Tomás JM, Bengoechea JA. Capsule polysaccharide is a bacterial decoy for antimicrobial peptides. Microbiology. 2008;154(12):3877–86.

45. Campos MA, Vargas MA, Regueiro V, Llompart CM, Albertí S, Bengoechea JA. Capsule polysaccharide mediates bacterial resistance to antimicrobial peptides. Infect Immun. 2004;72(12):7107–14.

46. Needham BD, Trent MS. Fortifying the barrier: the impact of lipid A remodelling on bacterial pathogenesis. Nat Rev Microbiol. 2013;11(7):467–81.

47. Gruenheid S, Le Moual H. Resistance to antimicrobial peptides in Gram-negative bacteria. FEMS Microbiol Lett. 2012 May;330(2):81–9.

48. Maeshima N, Fernandez RC. Recognition of lipid A variants by the TLR4-MD-2 receptor complex. Front Cell Infect Microbiol. 2013;3(February):3.

49. Raetz CR, Reynolds MC, Trent SM, Bishop RE. Lipid A modification in Gram-negative bacteria. Annu Rev Biochem. 2007;76(3):295–329.

50. Doorduijn DJ, Rooijakkers SHM, van Schaik W, Bardoel BW. Complement resistance mechanisms of *Klebsiella pneumoniae*. Immunobiology. 2016;221(10):1102–9.

51. Matsuura M. Structural modifications of bacterial lipopolysaccharide that facilitate Gram-negative bacteria evasion of host innate immunity. Front Immunol. 2013;4:109.

52. Andrews JM. Determination of minimum inhibitory concentrations. J Antimicrob Chemother. 2001 Jul;48:5–16.

53. Satlin MJ, Weinstein MP, Patel J, Romney M, Kahlmeter G, Giske CG, et al. Clinical and Laboratory Standards Institute (CLSI) and European Committee on Antimicrobial Susceptibility Testing (EUCAST) position statements on polymyxin B and colistin clinical breakpoints. Clin Infect Dis. 2020;

54. Loman NJ, Quinlan AR. Poretools: a toolkit for analyzing nanopore sequence data. Bioinformatics. 2014;30(23):3399–401.

55. van Mansfeld R, de Been M, Paganelli F, Yang L, Bonten M, Willems R. Within-host evolution of the Dutch high-prevalent *Pseudomonas aeruginosa* clone ST406 during chronic colonization of a patient with cystic fibrosis. PLoS One. 2016;11(6):e0158106.

56. Bankevich A, Nurk S, Antipov D, Gurevich AA, Dvorkin M, Kulikov AS, et al. SPAdes: a new genome assembly algorithm and its applications to single-cell sequencing. J Comput Biol. 2012 May;19(5):455–77.

57. Seemann T. Prokka: rapid prokaryotic genome annotation. Bioinformatics. 2014;30(14):2068–9.

58. Zankari E, Hasman H, Cosentino S, Vestergaard M, Rasmussen S, Lund O, et al. Identification of acquired antimicrobial resistance genes. J Antimicrob Chemother. 2012;67:2640–4.

59. Langmead B, Salzberg SL. Fast gapped-read alignment with Bowtie 2. Nat Methods. 2012 Apr;9(4):357–9.

60. Li H, Handsaker B, Wysoker A, Fennell T, Ruan J, Homer N, et al. The Sequence Alignment/Map format and SAMtools. Bioinformatics. 2009 Aug 15;25(16):2078–9.

61. Siguier P, Perochon J, Lestrade L, Mahillon J, Chandler M. ISfinder: the reference centre for bacterial insertion sequences. Nucleic Acids Res. 2006;34:D32–6.

62. Hawkey J, Hamidian M, Wick RR, Edwards DJ, Billman-Jacobe H, Hall RM, et al. ISMapper: Identifying transposase insertion sites in bacterial genomes from short read sequence data. BMC Genomics. 2015;16(1):667.

63. Briskine R V., Shimizu KK. Positional bias in variant calls against draft reference assemblies. BMC Genomics. 2017;18(1):263.

64. Llobet E, Martínez-Moliner V, Moranta D, Dahlström KM, Regueiro V, Tomás A, et al. Deciphering tissue-induced *Klebsiella pneumoniae* lipid A structure. Proc Natl Acad Sci U S A. 2015 Nov 2;112(46):E6369–E6378.

65. Llobet E, Campos MA, Giménez P, Moranta D, Bengoechea JA. Analysis of the networks controlling the antimicrobial-peptide-dependent induction of *Klebsiella pneumoniae* virulence factors. Infect Immun. 2011;79(9):3718–32.

66. El Hamidi A, Tirsoaga A, Novikov A, Hussein A, Caroff M. Microextraction of bacterial lipid A: easy and rapid method for mass spectrometric characterization. J Lipid Res. 2005;46(8):1773–8.

67. Napier BA, Burd EM, Satola SW, Cagle SM, Ray SM, McGann P, et al. Clinical use of colistin induces cross-resistance to host antimicrobials in *Acinetobacter baumannii*. MBio. 2013;4(3):e00021–13.

68. Brenner S. The genetics of *Caenorhabditis elegans*. Genetics. 1974;77(1):95–104.

69. Kurz CL, Chauvet S, Andrès E, Aurouze M, Vallet I, Michel GPF, et al. Virulence factors of the human opportunistic pathogen *Serratia marcescens* identified by *in vivo* screening. EMBO J. 2003 Apr 1;22(7):1451–60.

70. Porta-de-la-Riva M, Fontrodona L, Villanueva A, Cerón J. Basic *Caenorhabditis elegans* methods: synchronization and observation. J Vis Exp. 2012;64(64):e4019.

71. Jeannot K, Bolard A, Plesiat P. Resistance to polymyxins in Gram-negative organisms. Int J Antimicrob Agents. 2017;49(5):526–35.

72. Olaitan AO, Diene SM, Kempf M, Berrazeg M, Bakour S, Gupta SK, et al. Worldwide emergence of colistin resistance in *Klebsiella pneumoniae* from healthy humans and patients in Lao PDR, Thailand, Israel, Nigeria and France owing to inactivation of the PhoP/PhoQ regulator *mgrB*: an epidemiological and molecular study. Int J Antimicrob Agents. 2014;44(6):500–7.

73. Cheng Y-H, Lin T-L, Lin Y-T, Wang J-T. Amino acid substitutions of CrrB responsible for resistance to colistin through CrrC in *Klebsiella pneumoniae*. Antimicrob Agents Chemother. 2016;60(6):3709–16.

74. Simpson BW, Trent MS. Pushing the envelope: LPS modifications and their consequences. Nat Rev Microbiol. 2019;17:403–16.

75. Li J, Nation RL, Kaye KS. Polymyxin antibiotics: from laboratory bench to bedside. 2019.

76. Schnetz K, Rak B. IS5: a mobile enhancer of transcription in *Escherichia coli*. Proc Natl Acad Sci. 1992;89:1244–8.

77. Aghapour Z, Gholizadeh P, Ganbarov K, Bialvaei AZ, Mahmood SS, Tanomand A, et al. Molecular mechanisms related to colistin resistance in Enterobacteriaceae. Infect Drug Resist. 2019;12:965–75.

78. Nielsen LE, Snesrud EC, Onmus-Leone F, Kwak YI, Avilés R, Steele ED, et al. IS*5* element integration, a novel mechanism for rapid *in vivo* emergence of tigecycline nonsusceptibility in *Klebsiella pneumoniae*. Antimicrob Agents Chemother. 2014;58(10):6151–6.

79. Whitfield C. Biosynthesis and assembly of capsular polysaccharides. Annu Rev Biochem. 2006;75:39–68.

80. Lacour S, Bechet E, Cozzone AJ, Mijakovic I, Grangeasse C. Tyrosine phosphorylation of the UDP-glucose dehydrogenase of *Escherichia coli* is at the crossroads of colanic acid synthesis and polymyxin resistance. PLoS One. 2008;3(8):e3053.

81. Obadia B, Lacour S, Doublet P, Baubichon-Cortay H, Cozzone AJ, Grangeasse C. Influence of tyrosine-kinase Wzc activity on colanic acid production in *Escherichia coli* K12 cells. J Mol Biol. 2007;367(1):42–53.

82. Grangeasse C, Obadia B, Mijakovic I, Deutscher J, Cozzone AJ, Doublet P. Autophosphorylation of the *Escherichia coli* protein kinase Wzc regulates tyrosine phosphorylation of Ugd, a UDP-glucose dehydrogenase. J Biol Chem. 2003;278(41):39323–9.

83. Pal S, Verma J, Mallick S, Rastogi SK, Kumar A, Ghosh AS. Absence of the glycosyltransferase WcaJ in *Klebsiella pneumoniae* ATCC13883 affects biofilm formation, increases polymyxin resistance and reduces murine macrophage activation. Microbiology. 2019;165(8):891–904.

84. Ciampi MS. Rho-dependent terminators and transcription termination. Microbiology. 2006;152(9):2515–28.

85. Jayol A, Poirel L, Brink A, Villegas M-V, Yilmaz M, Nordmann P. Resistance to colistin associated with a single amino acid change in protein PmrB among *Klebsiella pneumoniae* isolates of worldwide origin. Antimicrob Agents Chemother. 2014 Aug;58(8):4762–6.

86. Cheng Y-H, Lin T-L, Pan Y-J, Wang Y-P, Lin Y-T, Wang J-T. Colistin-resistant mechanisms of *Klebsiella pneumoniae* in Taiwan. Antimicrob Agents Chemother. 2015 Feb 17;59(5):2909–13.

87. Choi MJ, Ko KS. Mutant prevention concentrations of colistin for *Acinetobacter baumannii*, *Pseudomonas aeruginosa* and *Klebsiella pneumoniae* clinical isolates. J Antimicrob Chemother. 2014;69(1):275–7.

88. Cheong HS, Kim SY, Wi YM, Peck KR, Ko KS. Colistin heteroresistance in *Klebsiella pneumoniae* isolates and diverse mutations of PmrAB and PhoPQ in resistant subpopulations. J Clin Med. 2019;8(9):1444.

89. Mills G, Dumigan A, Kidd T, Hobley L, Bengoechea JA. Identification and characterization of two *Klebsiella pneumoniae lpxL* lipid A late acylstransferases and their role in virulence. Infect Immun. 2017;85(9):e00068–17.

90. Brochado AR, Telzerow A, Bobonis J, Banzhaf M, Mateus A, Selkrig J, et al. Species-specific activity of antibacterial drug combinations. Nature. 2018;559(7713):259–63.

91. Imamovic L, Sommer MOA. Use of collateral sensitivity networks to design drug cycling protocols that avoid resistance development. Sci Transl Med. 2013;5(204):204ra132.

92. Imamovic L, Ellabaan MMH, Dantas Machado AM, Citterio L, Wulff T, Molin S, et al. Drug-driven phenotypic convergence supports rational treatment strategies of chronic infections. Cell. 2018;172(1-2):121–134.e14.

93. Gotoh Y, Eguchi Y, Watanabe T, Okamoto S, Doi A, Utsumi R. Two-component signal transduction as potential drug targets in pathogenic bacteria. Curr Opin Microbiol. 2010;13(2):232–9.

