## Supplemental Table S1 for "*In vitro* evolution of colistin resistance in the *Klebsiella pneumoniae* complex follows multiple evolutionary trajectories with variable effects on fitness and virulence characteristics"

| **Primer Name** | **Sequence (5'‑3')** |
| --- | --- |
| KP209 IS903B plasmid1 124Kbp F | CCT GGA TGA GAG GTA GAT GG |
| KP209 IS903B plasmid1 124Kbp R | GAA GCG TCG TAT CCC ATA AC |
| KP209 ISKpn1 chromosome 4.6Mbp F | TGC TGG TGT CGT TTA TCT GC |
| KP209 ISKpn1 chromosome 4.6Mbp R | ATG GAC AGA TAT CCG CGT TC |
| KP209 IS26 plasmid1 127Kbp F | GGC AGT GCA AAT CGA AAA AT |
| KP209 IS26 plasmid1 127Kbp R | TCA TTC AGC ACG CAA AAT TC |
| KP209 ISKpn1 chromosome 533Kbp F | ATG TCG ATC CGA TTC GTC A |
| KP209 ISKpn1 chromosome 533Kbp R | GTT CGT CAG CAC CCA TGT TT |
| KP040 ISKpn38 plasmid1 302Kbp F | TCA GGA AAA AGC GAG TCG TT |
| KP040 ISKpn38 plasmid1 302Kbp R | TTC GTT GAG CGT ATC GAG TG |
| KP040 IS1A plasmid1 45Kbp F | CAT GCA GCA TTG GTA CAA CC |
| KP040 IS1A plasmid1 45Kbp R | ACG TTT TCA ACC CAC AGA GG |
| KP040 ISEhe3 chromosome 1.7Mbp F | TGG AGC TGG TAT ACC GGT TC |
| KP040 ISEhe3 chromosome 1.7Mbp R | CGA CCA GGT GGC AAG TTT AT |
| KP040 IS102 chromosome 437Kbp F | GTC AGT ATT GGA CCA GAT CGT |
| KP040 IS102 chromosome 437Kbp R | TGC AAG CAG ACC TTT ATC AAA AT |
| KP040 IS5 chromosome 950Kbp F | GAT TGA TTG CCT GCT CAC CG |
| KP040 IS5 chromosome 950Kbp R | ACC AGA CCG AAT GTT ATT GCA |
| KP257 IS903B plasmid2 36.9Kbp F | G­­AA GCC ATG CTG GAT AAG GA |
| KP257 IS903B plasmid2 36.9Kbp R | GGT CTT AAT GCC CAG CAC AT |
| KP257 IS102 plasmid1 ­146Kbp F | ATA TCG CTG AAC CAG TGC TC |
| KP257 IS102 plasmid1 146Kbp R | ACA TCG AGT GGC TTC TGA AA |
| KP257 IS102 plasmid2 28Kbp F | TGG GTT ACC ACC AAA CGA AT |
| KP257 IS102 plasmid2 28Kbp R | GCT TTC AGA GCC TGG ATG AC |
