## Supplemental Table S2 for "*In vitro* evolution of colistin resistance in the *Klebsiella pneumoniae* complex follows multiple evolutionary trajectories with variable effects on fitness and virulence characteristics"

| **Strain** | **Sample** | **Type** | **Size of genome assembly** | **Number of scaffolds** | **Illumina reads mapped to reference** | **Average coverage of mapped Illumina reads** |
| --- | --- | --- | --- | --- | --- | --- |
| **KP209** | Colistin‑susceptible | Axenic | 5463023 bp | 1 Chromosome | 4976524 | 224­.3 |
|  |  |  |  | 5 plasmids |  |  |
|  | Colistin‑resistant | Axenic | NA | NA | 4310150 | 192.7 |
|  | Day 1 | Population | NA | NA | 5273935 | 226.1 |
|  | Day 2 | Population | NA | NA | 3498310 | 148.9 |
|  | Day 3 | Population | NA | NA | 6034443 | 265.1 |
|  | Day 4 | Population | NA | NA | 8381010 | 360.5 |
|  | Day 5 | Population | NA | NA | 10284032 | 439.1 |
| **KP040** | Colistin‑susceptible | Axenic | 5970459 bp | 1 chromosome | 4645736 | 190.5 |
|  |  |  |  | 1 plasmid |  |  |
|  | Colistin‑resistant | Axenic | NA | NA | 1387900 | 56.9 |
|  | Day 1 | Population | NA | NA | 2389602 | 96.2 |
|  | Day 2 | Population | NA | NA | 2145375 | 83.7 |
|  | Day 3 | Population | NA | NA | 3207128 | 125.2 |
|  | Day 4 | Population | NA | NA | 3232802 | 124 |
|  | Day 5 | Population | NA | NA | 2556849 | 96.3 |
|  | Day 6 | Population | NA | NA | 3835645 | 148.7 |
|  | Day 7 | Population | NA | NA | 2807670 | 105.6 |
| **KP257** | Colistin‑susceptible | Axenic | 5557780 bp | 1 chromosome | 2688902 | 119.1 |
|  |  |  |  | 3 plasmids |  |  |
|  | Colistin‑resistant | Axenic | NA | NA | 2073063 | 91.1 |
|  | Day 1 | Population | NA | NA | 4120472 | 172.3 |
|  | Day 2 | Population | NA | NA | 3878524 | 161.7 |
|  | Day 3 | Population | NA | NA | 4176568 | 173.4 |
|  | Day 4 | Population | NA | NA | 4669134 | 194.3 |
|  | Day 5 | Population | NA | NA | 3334800 | 139.7 |
|  | Day 6 | Population | NA | NA | 4707153 | 193.4 |
|  | Day 7 | Population | NA | NA | 3884435 | 160.5 |
| **KV402** | Colistin‑susceptible | Axenic | 5629276 bp | 1 chromosome | 2110840 | 92.9 |
|  |  |  |  | 2 plasmids |  |  |
|  | Colistin‑resistant | Axenic | NA | NA | 1453643 | 63.9 |
|  | Day 1 | Population | NA | NA | 6070185 | 255.2 |
|  | Day 2 | Population | NA | NA | 4501469 | 186.2 |
|  | Day 3 | Population | NA | NA | 5664425 | 234.4 |
|  | Day 4 | Population | NA | NA | 6186379 | 256.1 |
|  | Day 5 | Population | NA | NA | 5829343 | 241.2 |
|  | Day 6 | Population | NA | NA | 6218915 | 254.1 |
