## Supplemental Table S3 for "*In vitro* evolution of colistin resistance in the *Klebsiella pneumoniae* complex follows multiple evolutionary trajectories with variable effects on fitness and virulence characteristics"

| **Strain** | **Associated feature(s)** | **Location** | **Scaffold** | **Reference sequence** | **Evolved strain** | **Amino acid change** |
| --- | --- | --- | --- | --- | --- | --- |
| **KP209** | *phoQ* | 4336315 | Chromosome | G GGC GCA GCG TGA | G | 97-WAQRN 🡺97-C |
|  | *phoQ* | 4336791 | Chromosome | C | A | R256S |
|  | *pmrB* | 4692375 | Chromosome | T | A | Y209N |
| **KP040** | *yejM* | 304945 | Chromosome | G | C | H20Q |
|  | Intergenic SNP: promoter region *ecpR,* promoter region *phnC* | 2954735 | Chromosome | T | G | *Not applicable* |
|  | *rho* | 3907646 | Chromosome | C | C GCG ATG TTC TGC | 189-QS 🡺 189QNIAQS |
| **KP257** | *nlhH* | 175249 | Chromosome | C | CCA | 155-GGHLALVTALRLK… 🡺 155-GQGISPWSRLCA-**STOP** |
|  | Intergenic SNP: promoter region *yedY,* promoter region *csrD* | 720258 | Chromosome | G | A | *Not applicable* |
|  | *yxeP* | 2443607 | Chromosome | C | T | A384V |
|  | *yxeP* | 2443610 | Chromosome | C | G | A385G |
|  | *phoQ* | 3417233 | Chromosome | G | A | G385S |
|  | *afR* | 4155563 | Chromosome | G | C | P12R |
|  | *lptD* | 4659739 | Chromosome | C | G | S646R |
| **KV402** | *wcaJ* | 865365 | Chromosome | A CAA CAT CTT TAA T | A | 222-KIKDV… 🡺 222-KLDN-**STOP** |
|  | *phoQ* | 4702372 | Chromosome | A | T | L210Q |
|  | *phoP* | 4703101 | Chromosome | C | T | D191N |
|  | Intergenic SNP: promoter region hyp. protein, terminator region *traM* | 13871 | Plasmid | A | C | *Not applicable* |
