## Supplemental Table S4 for "*In vitro* evolution of colistin resistance in the *Klebsiella pneumoniae* complex follows multiple evolutionary trajectories with variable effects on fitness and virulence characteristics"

### Strain KP209

|  | **Location** | |  |  |
| --- | --- | --- | --- | --- |
| **IS element** | **Chromosome** | **Plasmid 1** | **ISMapper result** | **PCR Results** |
| **IS*1351*** | 2273711 - 2274917 | Not detected | Stable | NA |
| **IS*1400*** | 409379 - 408188 | Not detected | Stable | NA |
| **IS*26*** | Not detected | 3039 - 3858 | Stable | NA |
|  |  | 46202 - 47021 | Stable | NA |
|  |  | 103693 - 102874 | Stable | NA |
|  |  | 127823 - 128642 | Exision | No changes in integration |
| **IS*903B*** | Not detected | 124785 - 125840 | Integration | No changes in integration |
| **IS*Ec52*** | Not detected | 90987 - 92234 | Stable | NA |
| **IS*Kpn1*** | 533627 - 535071 | Not detected | Integration | No changes in integration |
|  | 1298963 - 1300407 |  | Stable | NA |
|  | 2693035 - 2691591 |  | Stable | NA |
|  | 3576388 - 3574944 |  | Stable | NA |
|  | 4229036 - 4227592 |  | Stable | NA |
|  | 4340825 - 4339381 |  | Stable | NA |
|  | 4684464 - 4683020 |  | Exision | No changes in integration |
|  | 4937819 - 4936375 |  | Stable | NA |
| **IS*Kpn26*** | Not detected | 29354 - 28159 | Stable | NA |
|  |  | 51021 - 49826 | Stable | NA |
| **IS*Sba14*** | Not detected | 77076 - 73370 | Stable | NA |

### Strain KP040

|  | **Location** | | | | | |  |  |
| --- | --- | --- | --- | --- | --- | --- | --- | --- |
| **IS element** | **Chromosome** | **Plasmid 1** | **Plasmid 2** | **Plasmid 3** | **Plasmid 4** | **Plasmid 5** | **ISMapper result** | **PCR Results** |
| **IS*102*** | 437037 |  | Not detected | Not detected | Not detected | Not detected | Integration | Novel integration on day 5 |
|  |  | 42230 - 41183 |  |  |  |  | Stable | NA |
|  |  | 203355 - 204410 |  |  |  |  | Stable | NA |
|  |  | 236546 - 237601 |  |  |  |  | Stable | NA |
| **IS*186B*** | 562310 - 563652 |  | Not detected | Not detected | Not detected | Not detected | Stable | NA |
|  |  | 106724 - 108062 |  |  |  |  | Stable | NA |
| **IS*1A*** | Not detected | 45702 - 44935 | Not detected | Not detected | Not detected | Not detected | Exision | No changes in integration |
| **IS*1B*** | Not detected | 147523 - 148290 | Not detected | Not detected | Not detected | Not detected | Stable | NA |
| **IS*1F*** | Not detected | 185253 - 186020 | Not detected | Not detected | Not detected | Not detected | Stable | NA |
| **IS*1R*** | 3445567 - 3444800 | Not detected | Not detected |  | Not detected | Not detected | Stable | NA |
|  | 3444800 - 3445567 |  |  |  |  |  | Stable | NA |
|  |  |  |  | 96541 - 97308 |  |  | Stable | NA |
| **IS*1X3*** | Not detected | 249803 - 249091 | Not detected | Not detected |  |  | Stable | NA |
| **IS*26*** | Not detected | 269007 - 269826 | Not detected |  | Not detected | Not detected | Stable | NA |
|  |  | 275149 - 275968 |  |  |  |  | Stable | NA |
|  |  |  |  | 92060 - 92879 |  |  | Stable | NA |
|  |  |  |  | 102074 - 102893 |  |  | Stable | NA |
| **IS*4321R*** | Not detected | 214068 - 215394 | Not detected | Not detected | Not detected | Not detected | Stable | NA |
|  |  | 236371 - 235045 |  |  |  |  | Stable | NA |
| **IS*5*** | 535139 - 536333 | Not detected | Not detected | Not detected | Not detected | Not detected | Stable | NA |
|  | 950081 |  |  |  |  |  | Integration | Novel integration on day 1 |
| **IS*903B*** | Not detected | 97940 - 96884 | Not detected |  | Not detected | Not detected | Stable | NA |
|  |  | 168847 - 167793 |  |  |  |  | Stable | NA |
|  |  | 170660 - 171716 |  |  |  |  | Stable | NA |
|  |  |  |  | 17990 - 19038 |  |  | Stable | NA |
| **IS*Cro1*** | Not detected | 294069 - 296767 | Not detected | Not detected | Not detected | Not detected | Stable | NA |
| **IS*Ec21*** | Not detected | 49613 - 50986 | Not detected | Not detected | Not detected | Not detected | Stable | NA |
| **IS*Ehe3*** | 1242490 - 1243718 | Not detected | Not detected | Not detected | Not detected | Not detected | Stable | NA |
|  | 1694238 - 1695466 |  |  |  |  |  | Stable | NA |
|  | 1753714 - 1754942 |  |  |  |  |  | Exision | No changes in integration |
|  | 3626554 - 3627782 |  |  |  |  |  | Stable | NA |
| **IS*Kpn14*** | Not detected | 160642 - 161409 | Not detected | Not detected | Not detected | Not detected | Stable | NA |
| **IS*Kpn1*** | 1860640 - 1859196 | Not detected | Not detected | Not detected | Not detected | Not detected | Stable | NA |
|  | 2518008 - 2519452 |  |  |  |  |  | Stable | NA |
|  | 2517676 - 2516232 |  |  |  |  |  | Stable | NA |
|  | 2577131 - 2578575 |  |  |  |  |  | Stable | NA |
|  | 2959264 - 2960708 |  |  |  |  |  | Stable | NA |
|  | 3692344 - 3693788 |  |  |  |  |  | Stable | NA |
|  | 4311386 - 4312830 |  |  |  |  |  | Stable | NA |
|  | 4878025 - 4876581 |  |  |  |  |  | Stable | NA |
| **IS*Kpn20*** | 611247 - 610052 |  | Not detected |  | Not detected | Not detected | Stable | NA |
|  | 706335 - 705140 |  |  |  |  |  | Stable | NA |
|  | 2969913 - 2971108 |  |  |  |  |  | Stable | NA |
|  | 4319319 - 4320514 |  |  |  |  |  | Stable | NA |
|  |  | 282044 - 283239 |  |  |  |  | Stable | NA |
|  |  |  |  | 7210 - 8405 |  |  | Stable | NA |
|  |  |  |  | 24339 - 25534 |  |  | Stable | NA |
| **IS*Kpn21*** | Not detected | 90829 - 93081 | Not detected | Not detected | Not detected | Not detected | Stable | NA |
|  |  | 302918 - 300666 |  |  |  |  | Stable | NA |
| **IS*Kpn24*** | Not detected | 30990 - 28537 | Not detected | Not detected | Not detected | Not detected | Stable | NA |
| **IS*Kpn26*** | 1611873 - 1613068 |  | Not detected | Not detected | Not detected | Not detected | Stable | NA |
|  | 2987253 - 2988448 |  |  |  |  |  | Stable | NA |
|  | 5228386 - 5229581 |  |  |  |  |  | Stable | NA |
|  |  | 43574 - 44769 |  |  |  |  | Stable | NA |
|  |  | 289171 - 290366 |  |  |  |  | Stable | NA |
| **IS*Kpn28*** | 1686614 - 1687707 |  | Not detected |  | Not detected | Not detected | Stable | NA |
|  | 1939240 - 1938147 |  |  |  |  |  | Stable | NA |
|  | 2802224 - 2803317 |  |  |  |  |  | Stable | NA |
|  | 4289154 - 4290247 |  |  |  |  |  | Stable | NA |
|  |  | 84936 - 86031 |  |  |  |  | Stable | NA |
|  |  | 105433 - 106528 |  |  |  |  | Stable | NA |
|  |  |  |  | 40456 - 41549 |  |  | Stable | NA |
| **IS*Kpn38*** | 1316492 - 1318089 |  | Not detected | Not detected |  | Not detected | Stable | NA |
|  |  | 87674 - 86077 |  |  |  |  | Stable | NA |
|  |  | 89232 - 90829 |  |  |  |  | Stable | NA |
|  |  | 302919 - 304516 |  |  |  |  | Integration | No changes in integration |
|  |  |  |  |  | 86969 - 88566 |  | Stable | NA |
| **IS*Ppu12*** | Not detected | 112459 - 115830 | Not detected | Not detected | Not detected | Not detected | Stable | NA |
| **IS*Sen3*** | Not detected | 175514 - 173559 | Not detected | Not detected | Not detected | Not detected | Stable | NA |
| **IS*Sen4*** | Not detected | 280209 - 278989 | Not detected | Not detected | Not detected | Not detected | Stable | NA |
|  |  | 292245 - 293465 |  |  |  |  | Stable | NA |

### Strain KP257

|  | **Location** | | | |  |  |
| --- | --- | --- | --- | --- | --- | --- |
| **IS element** | **Chromosome** | **Plasmid 1** | **Plasmid 2** | **Plasmid 3** | **ISMapper result** | **PCR Results** |
| **IS*102*** | Not detected | 146179‑147234 |  | Not detected | Exision | No changes in integration |
|  |  |  | 28056 ‑ 29111 |  | Exision | No changes in integration |
| **IS*1400*** | 4888362 ‑ 4887171 | Not detected | Not detected | Not detected | Stable | NA |
| **IS*1X3*** | Not detected | 133977 ‑ 134676 | Not detected | Not detected | Stable | NA |
| **IS*1X4*** | Not detected | Not detected | 17567 - 18334 | Not detected | Stable | NA |
| **IS*903B*** | Not detected | Not detected | 32740 - 33789 | Not detected | Stable | NA |
|  |  |  | 36985 - 38034 |  | Exision | No changes in integration |
| **IS*Ec21*** | Not detected | Not detected | 26032 - 27405 | Not detected | Stable | NA |
| **IS*Ehe3*** | Not detected | 8776 ‑ 7549 | Not detected | Not detected | Stable | NA |
| **IS*Eic1*** | Not detected | Not detected | 25486 - 24728 | Not detected | Stable | NA |
| **IS*Kpn1*** | 2465446 ‑ 2464002 | Not detected | Not detected | Not detected | Stable | NA |
|  | 4070871 ‑ 4069427 |  |  |  | Stable | NA |
|  | 4120943 ‑ 4122387 |  |  |  | Stable | NA |
|  | 4153758 ‑ 4152314 |  |  |  | Stable | NA |
| **IS*1S*** | Not detected | Not detected | 50676 ‑ 51438 | Not detected | Stable | NA |
| **IS*Kpn49*** | Not detected | 15667 ‑ 18106 | Not detected | Not detected | Stable | NA |

|  | **Location** | | |  |  |
| --- | --- | --- | --- | --- | --- |
| **IS element** | **Chromosome** | **Plasmid 1** | **Plasmid 2** | **ISMapper result** | **PCR Results** |
| **IS*1400*** | 5029253‑5030446 | Not detected | Not detected | Stable | NA |

**Strain KP402**
