## Supplemental Table S5 for "*In vitro* evolution of colistin resistance in the *Klebsiella pneumoniae* complex follows multiple evolutionary trajectories with variable effects on fitness and virulence characteristics"

| **Strain** | **Associated feature(s)** | **Location** | **Scaffold** | **Reference** | **Evolved strains** | **Amino acid change** |
| --- | --- | --- | --- | --- | --- | --- |
| **KP209** | *phoQ* | 4336315 | Chromosome | T GGG CGC AGC GTG | T | 97-WAQRN 🡺97-C |
|  | *phoQ* | 4336791 | Chromosome | C | A | R256S |
| **KP040** | *rho* | 3907646 | Chromosome | C | C GCG ATG TTC TGC | 189-QS 🡺 189-QNIAQS |
|  | *wzc* | 437037 | Chromosome | IS*5* insertion | | *Not applicable* |
|  | promoter region of *crrAB* and *crrC* | 950081 | Chromosome | IS*102* insertion | | *Not applicable* |
|  | Intergenic SNP: promoter region *yedY,* promoter region *csrD* | 720258 | Chromosome | G | A | *Not applicable* |
|  | *lptD* | 3246640 | Chromosome | TAAAA | TAAA | 764-ILPYQSSL-**STOP** 🡺  764-IYRTRAPCNAAGLQHHPHLRLIEMEKV-**STOP** |
| **KP257** | *phoQ* | 3417233 | Chromosome | G | A | G385S |
|  | *lptD* | 4659739 | Chromosome | C | G | S646R |
| **KV402** | *phoP* | 4703101 | Chromosome | C | T | D191N |
|  | *yciM* | 4950035 | Chromosome | T | G | V43G |
|  | *gcd* | 409843 | Chromosome | C | A | *Synonymous mutation* |
