## Supplementary figures and images for "*In vitro* evolution of colistin resistance in the *Klebsiella pneumoniae* complex follows multiple evolutionary trajectories with variable effects on fitness and virulence characteristics"

### Supplemental Figure S1

A

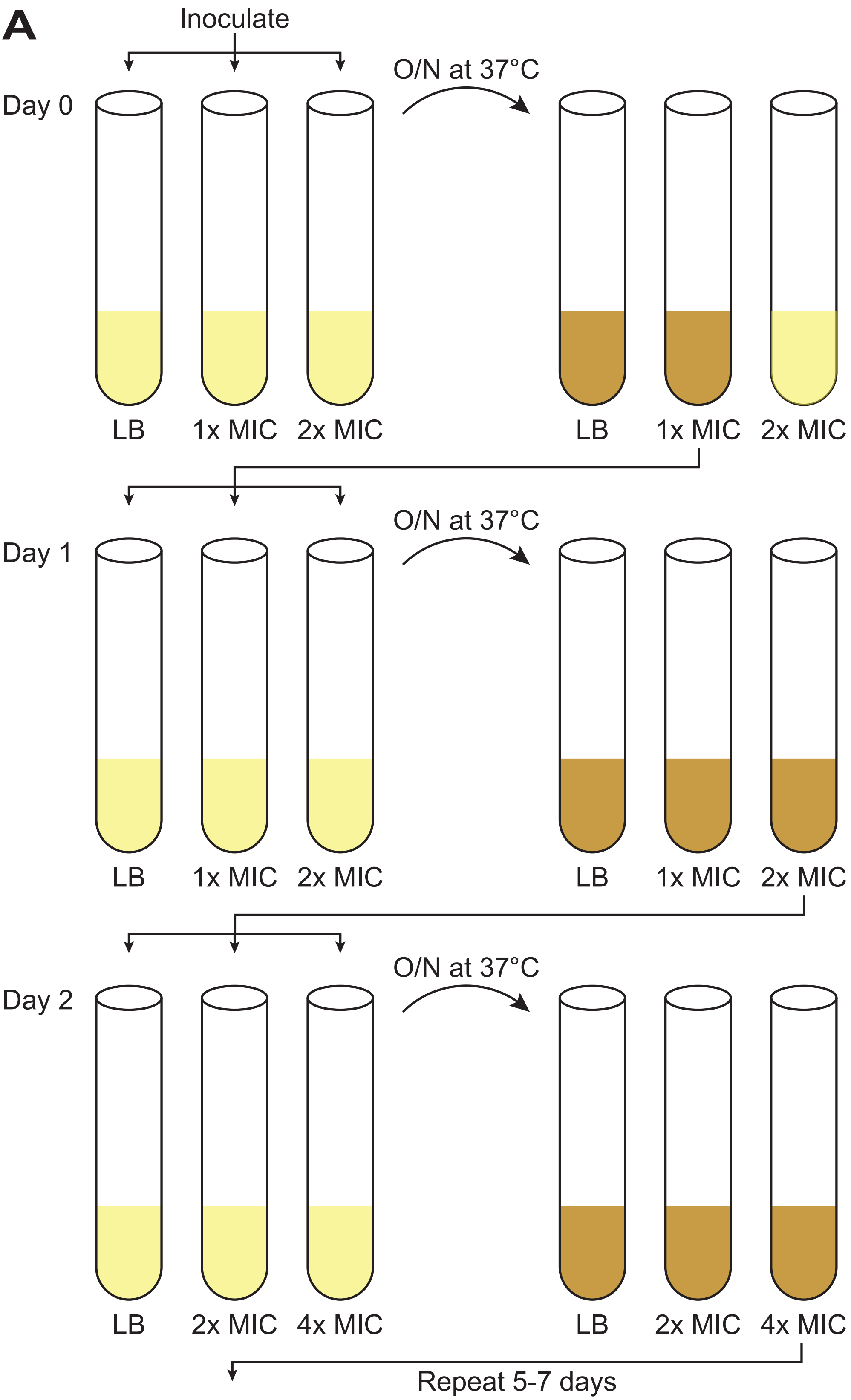


B

| Strain | KP209 | | KP040 | | KP257 | | KV402 | |
|---|---|---|---|---|---|---|---|---|
| MIC in LB (μg/ml) | 4 | | 0.5 | | 2 | | 2 | |
| Day 1 | 4 | 8 | 0.5 | 1 | 2 | 4 | 2 | 4 |
| Day 2 | 4 | 8 | 1 | 2 | 2 | 4 | 2 | 4 |
| Day 3 | 8 | 16 | 2 | 4 | 4 | 8 | 4 | 8 |
| Day 4 | 16 | 32 | 4 | 8 | 8 | 16 | 8 | 16 |
| Day 5 | 32 | 64 | 8 | 16 | 16 | 32 | 16 | 32 |
| Day 6 | 32 | 64 | 16 | 32 | 32 | 64 | 32 | 64 |
| Day 7 | 64 | 128 | 32 | 64 | 64 | 128 | 64 | 128 |

### Supplemental Figure S2

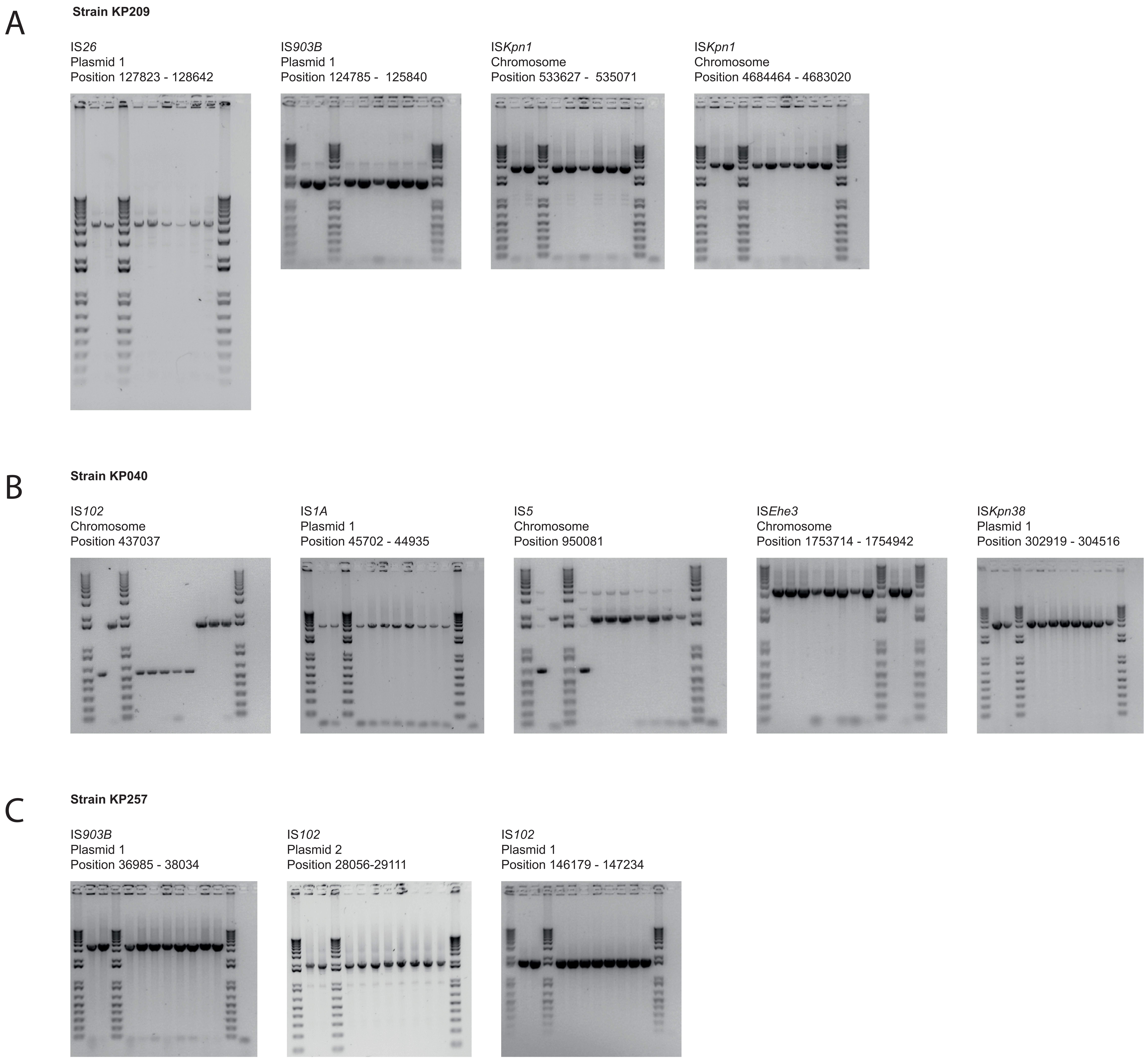
